# Capturing the nature of events and event context using Hierarchical Event Descriptors (HED)

**DOI:** 10.1101/2021.05.06.442841

**Authors:** Kay Robbins, Dung Truong, Stefan Appelhoff, Arnaud Delorme, Scott Makeig

**Author notes:** Corresponding author: Kay Robbins, Department of Computer Science University of Texas at San Antonio One UTSA Circle, San Antonio, TX 78249.

## Abstract

Event-related data analysis plays a central role in EEG and MEG (MEEG) and other neuroimaging modalities such as fMRI. Choices about which events to report and how to annotate their full natures significantly influence the value, reliability, and reproducibility of neuroimaging datasets for further analysis and meta- or `mega-analysis`. A powerful annotation strategy using the new third-generation formulation of the Hierarchical Event Descriptors (HED) framework and tools (*hedtags.org*) combines robust event description with details of experiment design and metadata in a human-readable as well as machine-actionable form, making event annotation relevant to the full range of neuroimaging and other time series data. This paper considers the event design and annotation process using as a case study the well-known multi-subject, multimodal dataset of Wakeman and Henson made available by its authors as a Brain Imaging Data Structure (BIDS) dataset (*bids.neuroimaging.io*). We propose a set of best practices and guidelines for event annotation integrated in a natural way into the BIDS metadata file architecture, examine the impact of event design decisions, and provide a working example of organizing events in MEEG and other neuroimaging data. We demonstrate how annotations using HED can document events occurring during neuroimaging experiments as well as their interrelationships, providing machine-actionable annotation enabling automated within- and across-experiment analysis and comparisons. We discuss the evolution of HED software tools and have made an accompanying HED-annotated BIDS-formatted edition of the MEEG data of the Wakeman and Henson dataset (*openneuro.org, ds003645*).

## 1. Introduction

EEG (electroencephalography) and MEG (magnetoencephalography) neuroimaging, collectively known as MEEG, are non-invasive brain imaging technologies for capturing neuroelectromagnetic brain dynamic records at millisecond-scale sampling rates. As MEEG records brain signals occurring on the time scale of individual thoughts and actions, event-related data analysis plays a central role in MEEG and other types of neuroimaging experiments. Because of the essential role that event markers and their annotations play in linking experimental data to the unfolding of the experiment, incomplete event reporting using event annotations that are inaccurate, overly simple, or absent represent significant barriers to analysis of shared neuroimaging data. But thoughtful choices as to how *events* are measured, identified, and annotated can greatly improve the utility of the collected data for both immediate and long-term analyses.

Good annotation tools and standards can also incorporate useful information about experimental design, participant tasks, data features (for example eyeblinks, movement artifact, ictal activity), and other metadata *into* the collected and later shared data, thereby making the data ready for efficient within- and across-study analyses using a variety of approaches. Though here we focus on MEEG applications, event annotation standards and practices essential for MEEG data analysis can be applied equally well to other types of neuroimaging time series data including fMRI. For example, growing appreciation of the importance of embodied cognition on mental life (Shapiro, 2019), new lightweight, low cost methods of recording details of brain activities and motor behavior of experiment participants (Casson, 2019) (Jas et al., 2021) (Vitali & Perkins, 2020), and emergence of the practice of recording both brain activity and behavior (as well as psychophysiology) at higher resolution in a broader range of tasks and task environments (often termed Mobile Brain/Body Imaging or MoBI) (Makeig et al., 2009), make development of a suitable and more comprehensive data annotation framework ever more urgent.

### Events

In everyday life, we use the term “*event*” to describe some experience (or sequence of interrelated experiences) unfolding through time that has some significance distinguishing it from other preceding, concurrent, and succeeding events. Events in this sense may be brief (e.g., the experience of hearing an unexpected click) or may unfold over any time period (e.g., the experience of viewing a movie, or of repeatedly performing a cognitive task during a neuroimaging experiment).

Moreover, experiences we may refer to as events may be nested in time. For example we may recall, as a meaningful event, our emotional response to viewing the surprising first clip of a particular scene in a movie presented to us during a neuroimaging recording session. However, we may equally well recall, and think of as an event, our experience of viewing that clip, or our experience of viewing the whole scene, or the whole movie ─ or, of participating in the entire recording session. In recounting another experienced ‘event,’ we typically recall and describe its critical transition points (e.g., “game kickoff”, “the final movie credits beginning to scroll”, “my feeling the moment after the electrode cap came off”). These we might liken to phase transition moments in a time-limited dynamic process.

### Event markers

In neuroimaging time-series recordings, metadata, experiment events are typically recorded using event markers marking that each mark the time of some phase transition or other point of interest in the unfolding event or event process (most often, time of onset). Unfortunately, in practice these event markers are often themselves labelled and referred to as “events”, risking conceptual confusion.

Each event marker designates a single time point, typically expressed as a time offset from the start of the time series recording. To be useful, the event marker must be associated with metadata that includes information about the type of event phase transition it marks, a reference to the ongoing event process it marks, as well as a description of the nature of that event. The description of the event is most conveniently associated with the event marker marking its onset. Event markers of later phase transitions in the event (e.g., its offset) need not repeat this description if they include an unequivocal reference to the event. As well as marking event onsets and offsets, event markers may mark other meaningful event phase transitions – for example the moments at which the trajectories of balls thrown by a participant in a juggling experiment reach their apex or a presented sound reaches maximum volume. Analyses aimed to better understand how brain activity supports skilled juggling or speech comprehension may well strongly benefit from identifying and then marking these moments in the experiment data record.

Fig. 1 illustrates these concepts schematically. During a task condition in which spatial target ‘+’ images are briefly presented at different screen positions; the participant is instructed to reach to touch the center of the current or most recently displayed target. HED annotations associated with the event markers provide essential linkage between the event processes and the measured data. Below, we also show how HED annotation can also capture the relation of events to the experiment design.

**Figure 1.**
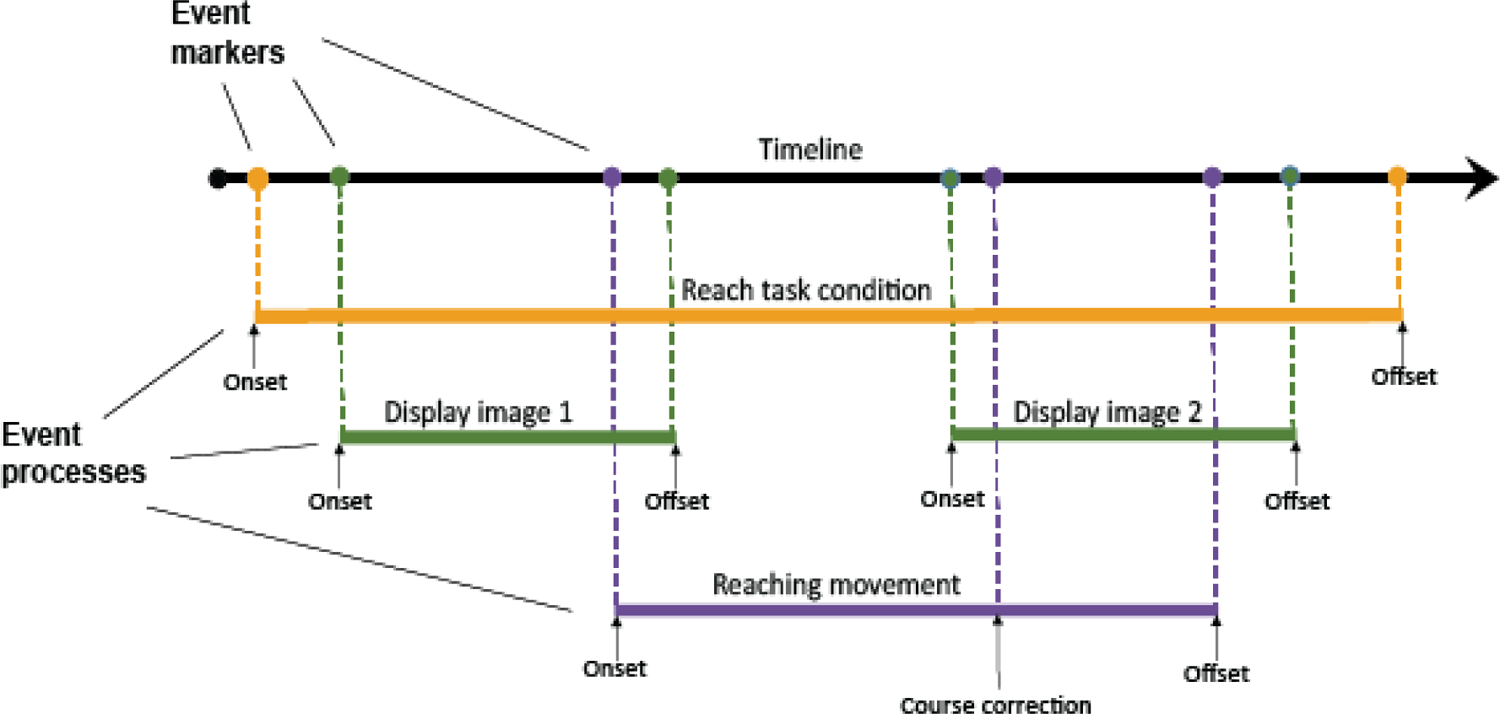
Schematic depiction of *event processes* during an MEEG experiment and their associated *event markers* displayed as dots on the timeline to index the latencies (time points) at which event process time boundaries or phase transitions occur in an experiment data recording. Below the (top, black) experiment timeline: (orange bar) *Onset* and *Offset* markers for reach task condition; (green bars) *Onset* and *Offset* markers of the visual presentation periods for image 1 and image 2 presentations; (purple bar) time course of a participant reach to touch movement. In addition to having *Onset* and *Offset* event markers, the reaching movement includes an intermediate marker of a recognized arm/hand trajectory course correction.

### Event context

An event occurring within longer-duration events (e.g., the experience of a stimulus presentation within a *supervening* task block in a neuroimaging session), and/or during temporally *overlapping* events, may be said to occur within the *context* of those events. Since event marker latencies use a common timeline, software tools may automatically add context information about other ongoing events (wholly concurrent or temporally overlapping) to the event marker metadata at their time of use in data search and analysis. In future, tools dealing with *event context* might be extended to facilitate desired analyses relating recorded brain dynamics to document the experienced *preceding* and/or anticipated *succeeding* events.

### Overview

This paper introduces a practical event design strategy and illustrates a set of best practices for event reporting and annotation based on combining the new third-generation formulation of the Hierarchical Event Descriptor (HED) annotation framework (Robbins et al., 2020) with the MEEG data storage architecture of the Brain Imaging Data Structure (BIDS) group (Gorgolewski et al., 2016) (Niso et al., 2018) (Pernet et al., 2019) (Holdgraf et al., 2019). The paper is organized around a case study using MEEG data from a publicly-available multi-participant, EEG/MEG and fMRI experiment by Daniel Wakeman and Richard Henson (Wakeman & Henson, 2015; abbreviated below as W-H) saved in conformity with the BIDS guidelines. The HED/BIDS integration of event annotation demonstrated and recommended here not only facilitates automatic and informative summarization of data; it also establishes a standardized interface for automated pipelines to search for, collect, and read and preprocess raw data, and perform automated event-related analysis using study-independent tools and vocabulary. In particular, the strategy enables analyses to be performed across stored datasets, even when these datasets do not have the same experiment design.

### W-H

The W-H experiment was conducted to develop methods for integrating multiple imaging modalities into analysis to increase the accuracy of functional and structural connectivity analyses. Nineteen participants completed two recording sessions spaced three months apart – one session recorded fMRI data (W-H-fMRI) and the other simultaneously recorded MEG and EEG data (W-H-MEEG). During each session, participants performed the same perceptual task, evaluating the symmetry of presented photographs of famous, unfamiliar, and scrambled faces. The participants pressed one of two keyboard keys with left or right index fingers, respectively, to indicate a subjective yes or no decision as to the relative spatial symmetry of each viewed image. The original, unannotated W-H dataset was made available on OpenNeuro (*openneuro.org*, *ds000117)*. Recently, we have shared a BIDS version of the W-H joint EEG/MEG data on OpenNeuro (*openneuro.org*, *ds003645*) with the more complete event organization and annotation discussed in this paper. Although we here focus on the MEEG portion of the W-H data set, the methods we demonstrate are equally applicable to annotation of fMRI or other neuroimaging time series data.

Unlike most MEEG experiments, the W-H overt face-symmetry judgment task was not itself of interest to the experimenters, who thus made no effort to judge whether participant responses had some objective basis in the face images themselves. Rather, the experiment was designed to covertly test recognition memory for the three types of face images. To this end, each individual face image was presented twice during the session. For half of the presented faces, the second presentation immediately followed the first. For the other half, the second presentation occurred after 5-15 intervening face image presentations. Famous faces were feature-matched to unfamiliar faces, and half the faces were female. Following the neuroimaging sessions, the authors also collected behavioral recognition memory performance measures from participants to allow testing for interactions between MEEG responses associated with individual image presentations and subsequent recognition memory for those images. These behavioral recognition memory data were also provided by the data authors for inclusion in our revised MEEG dataset.

Fig. 2 shows a schematic view of a typical event sequence in the W-H experiment. All of the session recordings were conducted using the same equipment, with the participant seated and facing a computer monitor throughout (top black timeline). The bottom two timelines show the introduced sensory events (visual screen image presentations, green timeline) and participant actions (left or right index finger button presses, purple timeline).

**Figure 2.**
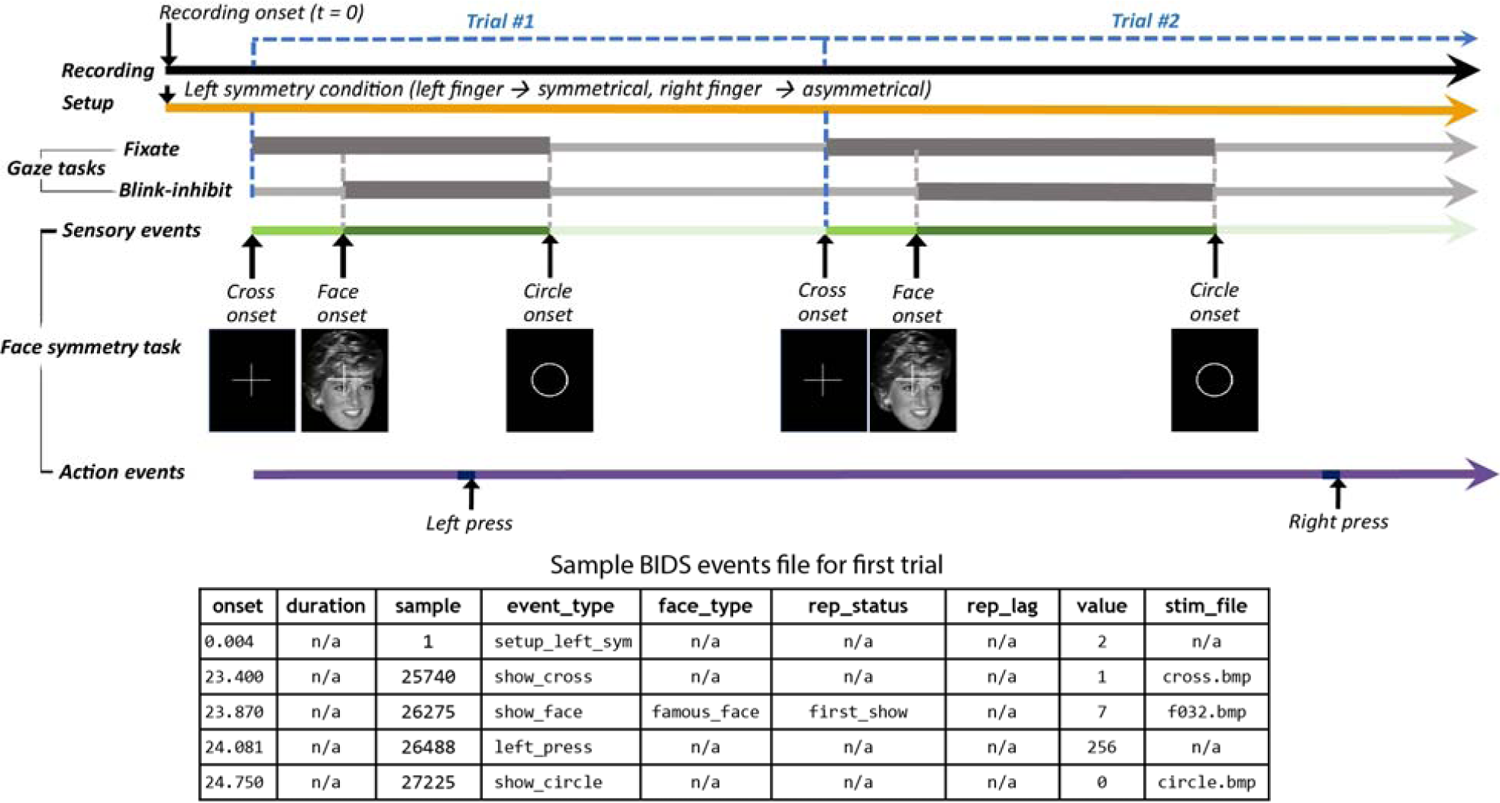
Schematic diagram of the temporal organization of events two trials of a W-H MEEG recording and excerpt of the BIDS task events file built using HED-based encoding strategies. Upper left: Recording begins. Recording setup includes selection of the key assignment for responses in the face symmetry judgment task. The participant was asked to fixate on a central cross, while presented, and to refrain from blinking while face images were presented. Lower timelines: Sensory events were visual image presentations; participant action events were key presses representing ‘face symmetry’ task responses. Bottom table: a BIDS task events file excerpt corresponding to the first trial in the data. We will use this example throughout the paper. (See an expanded version in Table 5, Section 3.1).

Some of the participants were instructed to follow each face image presentation onset with a left index finger key press to indicate above average facial symmetry and a right index finger press to indicate below average facial symmetry. The remaining participants used the opposite key assignment. The key assignment was in effect for all of the recordings associated with a particular participant (orange timeline). The participants were also instructed to fixate on the white cross and asked not to blink while the face was presented (thick gray gaze task timelines).

The fundamental problem addressed here is how to effectively describe events in a standardized form that is human-readable, machine-actionable, and analysis-ready – without placing undue burden on the annotator. The W-H-MEEG experiment has five regularly repeating types of events. We demonstrate how to create locally defined names (`show_cross, show_face, show_circle, left_press`, and `right_press`) using a standardized vocabulary (HED) and to associate these names with event markers, resulting in an analysis-ready annotated event stream.

The following section begins with a brief introduction to the HED system and, using the W-H MEEG experiment as a concrete example and explains the event annotation process, including annotations relating event types to the experiment design. Section 3 shows how these annotations can be organized within a BIDS dataset to achieve machine-actionable, analysis-ready annotation. Using the example developed in Sections 2 and 3, Section 4 examines the event design process and proposes a set of guidelines for effective design and annotation in neuroimaging research. We discuss what events should be reported, how the events should be encoded, and sketch planned further work to extend this encoding to include the relationship of the encoded events to participant task(s) and intent. Section 4 also summarizes the importance and potential impact of best-practice annotation strategies in making both stored and shared data more reproducible, interpretable and usable, first to the annotators themselves, then in any subsequent analysis enabled by effective data storage and sharing. We give a brief review and roadmap for future HED development in Section 5.

### 2. Machine-actionable event annotation using HED

The HED system is based on a collection of hierarchically organized terms (the base HED schema) that describe experiment events, condition variables, participant tasks, metadata, or the recording’s temporal structure. HED was specifically designed to encode information in a both human- and machine-actionable format to enable validation, search, identification, and analysis of events in neuroimaging or other time series datasets that include events with known timing.

The original HED implementation (first-generation) focused mainly on a description of stimuli and responses (Bigdely-Shamlo et al., 2013). The second-generation HED framework (Bigdely-Shamlo et al., 2016) included many vocabulary improvements, plus tools for validation, data search, and analysis and was accepted in 2019 as an optional standard for event annotation in BIDS formatted data.

HED has recently undergone an extensive third-generation redesign (HED-3G) to enable capture not only basic event and event marker descriptions, but also experimental conditions, temporal structure, and event context (Robbins et al., 2020). HED-3G provides a readily extensible basis for easily interpretable annotation of time series datasets for use in analysis, re- analysis, and shared data mega-analysis. HED-3G was officially released in August 2021 and is ready for widespread use in data archiving, sharing, analysis, and mega-analysis.

#### In this paper, we use the term HED to refer exclusively to HED-3G

The remainder of Section 2 works through the W-H-MEEG case study step-by-step to illustrate the HED annotation process and the major features of HED. The examples are organized so that the end result is a fully-annotated BIDS dataset.

### 2.1 A starting point for HED dataset event annotation

The HED base schema has seven top-level or root nodes as shown by the partially expanded schema tree in Fig. 3, left. The very basic HED event annotation shown in the table inset on the right is our starting point for development of comprehensive annotation.

**Figure 3.**
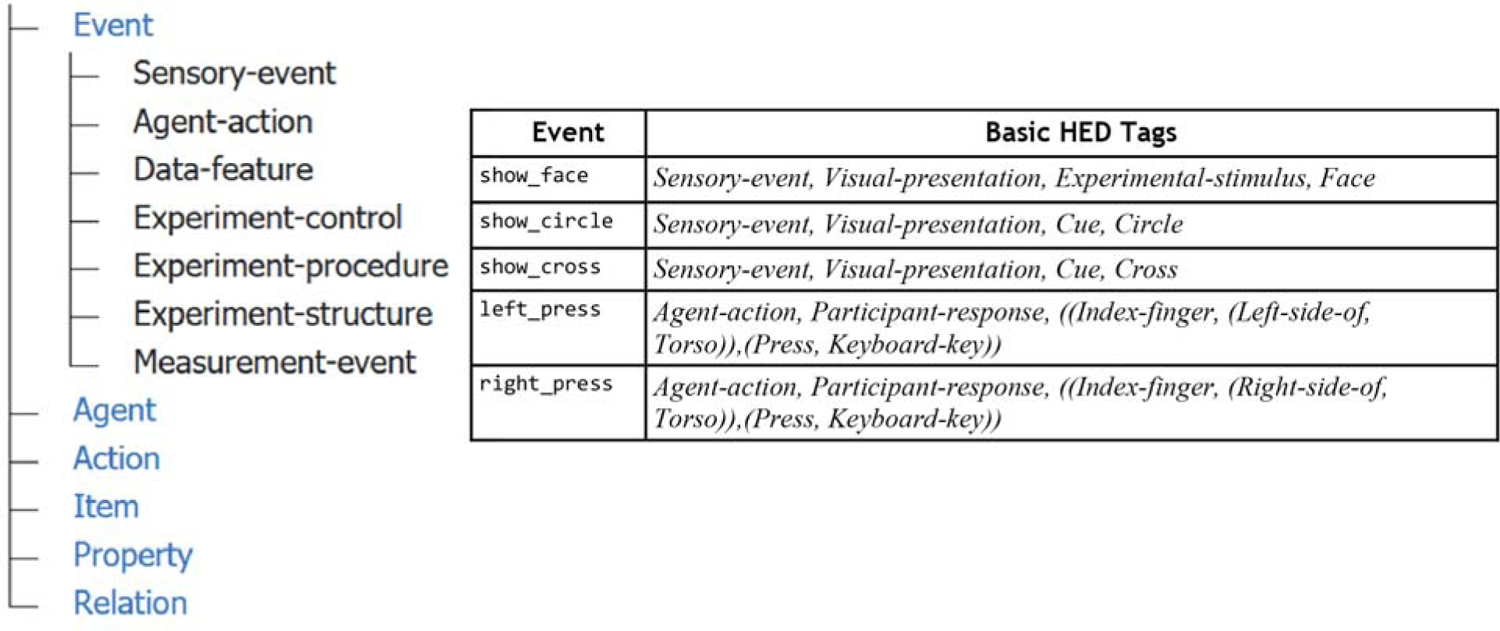
Left: Graphic of a partially expanded top-level HED schema tree (see www.hedtags.org/display_hed.html for a view of the complete schema in an easy-to-search expanding format). Right: A table with basic annotations of the five main W-H event marker types using HED. The left column of the table has user-defined terms used for convenience to refer to these event markers in the BIDS event files. The right column shows the underlying mapping of these terms to the common HED vocabulary.

To annotate events, users create comma-separated lists of terms selected from the HED base schema to describe the main events and concepts. This can be done as a table such as the one shown on the right in Fig. 3. Users first select an item from the *Event* top-level subtree to give a basic characterization of the event category (e.g., *Sensory-event*, *Agent-action*, *Data-feature*, etc.) for each of the main types of event markers. The top-level event categorization tag often serves as a primary search key for identifying events of interest. In addition to the event category, tags describing the sensory modality for sensory events or the type of action for agent actions are included next. In some sense, the annotation process can be thought of as using keywords from a structured vocabulary to tag events. The tag group *(Press, Keyboard-key)* in Fig. 3 then resembles a verb phrase, and the *(Index-finger, (Press, Keyboard-key))* tag group a sentence.

Additional tags should then be added to provide a more detailed description. For follow-on analyses, particularly comparisons of MEEG dynamics *across* experiments, having still more detailed annotation can add significant and enduring value to the data. In this example, adding annotations answering questions such as: “Which fingers pressed the keys?”, “How large were the cross, face image, and circle?”, “What colors were they?”, “Where were these images presented on the screen?”, and “For how long were they shown?”, can all add details to the annotation that could well prove of interest in further analyses and mega-analyses involving the data, even when (as here) the specific hypothesis testing for which the experiment was designed did not vary nor evaluate effects associated with answers to these questions.

While classical statistical testing assumes rigidly controlled experiments that involve controlled variation of at most a few features of interest, new statistical methods including machine learning can exploit diversity in labelled data to learn deep structure in the data – here, links between MEEG dynamics and human experience and behavior. In the past, the value of neuroimaging data for the researchers who created it chiefly depended primarily on the quality of the scientific paper they published using it. Increasingly, the value of neuroimaging data accruing to the data authors will also include the number and quality of further analyses that exploit the rich information contained in the dataset to power cross-study analysis.

### 2.2 Short and long form annotation

A critical usability innovation in third-generation HED is the requirement that each term in the HED schema must have a unique name (i.e., must only appear in one place in the schema). As a result, an annotator can tag using just a single end-node term (e.g., *Circle* in an *Item* hierarchy), rather than spelling out its full hierarchical schema path string (e.g., *Item/Object/Geometric-object/2D-shape/Ellipse/Circle*). Automated HED tools can then map such short-form tags to their complete (long-form) paths whenever the data are to be validated or analyzed. See the Tools section of the HED specification for links to tools written in Matlab, Python, and JavaScript to perform this mapping (*readthedocs.org/projects/hed-specification*).

The expanded long-form annotations allow tools collecting related events for analysis to find HED strings that belong to more general categories – for example, searching for event markers whose HED strings contain the more general term *2D-shape,* not only the more specific *Circle*. This type of organization is particularly useful for gathering data epochs time locked to a variety of events across datasets that have some feature or features in common, and/or have been annotated with different levels of detail.

The HED tag examples in this paper are given in short form for readability, and HED tags are always italicized. Supplementary Table 2 has examples of short form to long form tag expansion. While HED tags are case insensitive, by convention HED tags start with a capital letter and individual words in a tag are hyphenated. This convention makes it easier to pick out individual tags in a lengthy string of comma-separated tags. Also, HED tags cannot themselves contain blanks. In this paper we display locally-defined terms in fixed point type. Terms used in BIDS event files (e.g., `show_face` or `event_type`) use underbars as word separators to allow tools to directly map identifiers into program variables or structure fields.

### 2.3 Identifying event concepts using HED definitions

Fig. 3 (above) gives minimal HED annotations for the five most regularly occurring event types in the W-H dataset as described schematically in Fig. 2. This level of annotation allows analysts to isolate events of different types (stimulus events vs. participant actions, etc.), but does not provide sufficient detail to support advanced analysis and cross-study comparisons. Further, the annotation treats each event as occurring instantaneously, but the image presentation events have distinct onsets, durations, and offsets, all of which are known to affect brain dynamics measured by MEEG or fMRI.

**HED user event definitions** allow annotators to document the structure of the experiment, as laid out in Fig. 2, by “defining” or “declaring” experiment event-related concepts using names of their choosing and associating them with tag groups. During the annotation process, users can then use the defined names in place of the longer tag strings. HED definitions allow data authors to use shorthand terms from the colloquial lab jargon that they use in everyday lab conversations, while allowing data search and analysis to make use of the full HED annotations in analyses. Definitions also make it easier to initially identify and then later refine (all within the single definitions) annotations by adding tags to give further details. HED definitions thus can improve annotation process organization similar to the way first planning and then programming sub- functions can simplify the coding process and improve the resulting computer code.

Importantly, HED user definitions also play an integral role in assisting data authors in documenting experiment architecture, event temporal extent, and other dataset aspects. Consider a simple user definition (***Face-image***) for the presentation of a face image on a black background with a white fixation cross.

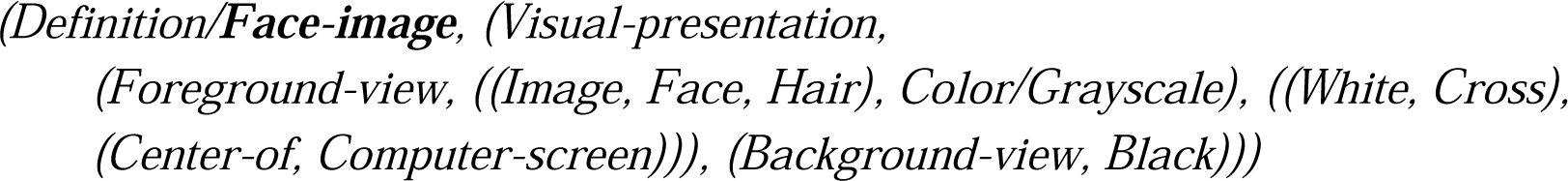

Here we embolden defined terms for ease of reading. For simplicity the definition uses short-form encoding (e.g., *Visual-presentation* instead of the full path string *Property/Sensory- property/Sensory-presentation/Visual-presentation*). Of course, this definition can be made more detailed, at any point in the annotation process. Note, however, that to avoid circularities HED definitions cannot be nested.

Once defined, annotators can use *Def/**Face-image*** in building annotations in place of the more complete but much longer and harder to remember tag string, thus increasing the readability of the dataset annotation while allowing the annotator to use (and more easily recall) terms that seem most natural to them.

Next, we focus on the use of HED definitions to annotate more of the temporal fine structure of the participant experience. The green timeline of Fig. 2 (Section 1) shows the time courses of the sensory events in the W-H data. The bright green bar marks the “pre-stimulus period” during which a white cross is displayed, while the dark green bar marks the time during which the face image is displayed, and the light green bar marks the period during which a white circle is displayed.

The boundaries between these displays are marked by the show_cross, show_face, and show_circle event markers, respectively. In the W-H experiment, face display ends when the circle image is presented. In addition, performance periods for two additional instructed eye-control tasks (represented by the thick gray timelines in Fig. 2) coincide with these events: 1) participants were asked to maintain eye gaze fixation on the white cross while it was displayed, and 2) to inhibit eye blinks during face image presentations.

Table 1 shows an expanded version of the table inset of Fig. 3 using definitions grouped with *Onset* and *Offset* tags to document temporal relationships between events indicated schematically in Fig. 2. (See Supplementary Table 1 for the complete annotation).

**Table 1.** The HED event marker annotations that capture repeating details of the W-H timeline

| Event | HED |
| --- | --- |
| `show_cross` | *Sensory-event, Cue, (Def/Cross-only, Onset), (Def/Fixation-task, Onset), (Def/Trial, Onset), (Def/Circle-only, Offset)* |
| `show_face` | *Sensory-event, Experimental-stimulus, (Def/Face-image, Onset), (Def/Blink-inhibition-task, Onset), (Def/Cross-only, Offset)* |
| `show_circle` | *Sensory-event, Cue, (Def/Circle-only, Onset), (Def/Trial, Offset), (Def/Face-image, Offset), (Def/Blink-inhibition-task, Offset), (Def/Fixation-task, Offset)* |
| `left_press` | *Agent-action, Participant-response, Def/Press-left-finger* |
| `right_press` | *Agent-action, Participant-response, Def/Press-right-finger* |

When a defined term such as *Face-image* is grouped with an *Onset* tag (e.g., such as *Def/Face-image* in the annotation for `show_face` in Table 1), the annotation represents the *Onset* marker of an event that unfolds over some duration. *Face-image* is assumed to be in effect until the next event in which a *Face-image* tag appears grouped with an *Onset* or *Offset* tag (the annotation for `show_circle` has an Offset). In the BIDS event file excerpt of Fig. 2 (Section 1), a `show_face` event onsets at time 23.87s, while the next `show_circle` event (whose annotation includes a *Def/**Face-image*** grouped with an *Offset* tag) occurs at 24.75s, Thus, the face image event presentation process unfolds over 24.75 − 23.87 = 0.88s.

Table 1 gives similar encodings for all the task-related sensory and participant action events. The ***Press-left-finger*** and ***Press-right-finger*** definitions of Table 1 do not include *Onset* or *Offset* tags because here only the time of key release was recorded; thus we only model these participant actions as instantaneous events that occur at a single moment in time.

### 2.4 Event context and temporal events

Effects of both *preceding and concurrent event context* on event-related MEEG brain dynamics have long been reported (Squires et al., 1977) although not frequently studied. When the full annotation of an event is assembled at time of data search or analysis, HED tools can automatically insert information about ongoing events in an *Event-context* tag group. For example, suppose a participant presses a key while a movie clip is playing. After creating a ***Play- movie*** definition to describe the movie presentation, the researcher can annotate the event marking the start of the movie with *(Def/Play-movie, Onset)* and the event marking the end of the movie with *(Def/Play-movie, Offset)*. HED tools can insert information that the movie was playing into the annotations of any concurrently occurring events. A future goal is to allow HED context tool annotation to also support studies of consequences of recent past events on the behavior and brain dynamics associated with current events.

### 2.5 Annotating experiment design and condition variables

The event marker sequences and the annotations described in the previous section define what happens during the experiment, but do not convey the purpose of the experiment or the relation of events to the underlying experimental design. A goal of HED is to provide convenient mechanisms for annotating this information in sufficient detail that tools can automatically extract and make use of experimental design information during analysis. HED supports the first steps in this process. This section introduces the *Condition-variable* tag and combines this tag with concept definitions to encode the W-H experimental design.

#### The W-H experiment design

The W-H experiment uses a factorial 3 × 3 matrix whose two factors are face type and repetition status, each with three levels. The primary author analyses (Henson et al., 2011) (Wakeman & Henson, 2015) (Henson et al., 2019) focused on face type analyses (with three levels corresponding to the display of famous, unfamiliar, and scrambled faces, respectively). The authors computed across-trial averaged event-related potentials (ERPs) and some frequency-based measures for MEEG responses to different types of face images with an underlying purpose of improving source localization by leveraging participant information obtained from multiple imaging modalities.

Each face (or scrambled face) image was shown twice during a session. The repetition status factor (with levels corresponding to the first display of an image, an immediately repeated display, and a delayed repeated display) encodes the position in the sequence of face image presentations with respect to their matching images. The delayed-repeat level indicates that the first presentation of this image occurred 5 to 15 face image presentations previously. The repetition status design variable was introduced to support study of the effects of image novelty and reinforcement on face recognition in the W-H data, supported by a later (behavior-only) face image recognition task session not included in the original version of the shared data.

### Documenting experiment control events

In BIDS datasets, information about changes in experiment conditions (e.g., in task or stimulus conditions) during a data recording session can be entered in one of two ways in the BIDS (`.tsv`) event file: either by inserting new columns in `.tsv` the event file table or by inserting new rows (events) in the events table.

Additional columns encode some item of information about every recorded event (row). The presence or absence of the informative condition is then indicated by the value in the cell of that column in every event row of the table ( used to indicate its absence or irrelevance). When n/a the information is relevant to only a small fraction of the recorded events, this can waste space and computation.

The alternative approach is to add new rows (event markers) to encode the information as their own events. Tools must then use context search to determine whether or not the information is relevant during the occurrence of any particular event. BIDS leaves the choice of representation (by row or column) to the user.

Table 2 summarizes the 3×3 W-H experimental design matrix and demonstrates how the experiment design can be encoded using HED. Here we will encode *design factor* information in *columns* added to the BIDS `.tsv` task events file. The factor names (see column 1 in Table 2) correspond to BIDS event-file column headings ( e_type and rep_status, respectively). The levels (s_face, unfamiliar_face scrambled_face) for the face type factor will appear as values in the face_type column of the BIDS event file. Similarly, the levels (_show first, immediate_repeat, delayed_repeat) of the repetition status factor appear as values in the rep_status column. The complete annotations are given in Supplementary Table 1.

**Table 2.** Encoding of the 3×3 experimental design for the W-H experiment using columns face_type and rep_status in the event files.

| Factor<br>(Column name) | Level<br>(Column value) | HED Annotation |
| --- | --- | --- |
| face_type | famous_face | **Description:** A face that should be recognized by the participants.<br>**HED:** (Definition/*Famous-face-cond*, (Condition-variable/*Face-type*, (Image, (Face, Famous)))) |
|  | unfamiliar_face | **Description:** A face that should not be recognized by the participants.<br>**HED:** (Definition/*Unfamiliar-face-cond*, (Condition-variable/*Face-type*, (Image, (Face, Unfamiliar)))) |
|  | scrambled_face | **Description:** A scrambled face image generated by the face 2D FFT.<br>**HED:** (Definition/*Scrambled-face-cond*, (Condition-variable/*Face-* |
|  |  | *type, (Image, (Face, Disordered))))* |
| rep_status | first_show | **Description:** Factor level indicating the first display of this face.<br>**HED:** (*Definition/First-show-cond, (Condition-variable/Repetition-status, Item-count/1)*) |
|  | immediate_repeat | **Description:** Factor level indicating this face was the same as previous.<br>**HED:** (*Definition/Immediate-repeat-cond, (Condition-variable/Repetition-status, Item-count/2, Item-interval/1)*) |
|  | delayed_repeat | **Description:** Factor level indicating face was seen 5 to 15 trials ago.<br>**HED:** (*Definition/Delayed-repeat-cond, (Condition-variable/Repetition-status, Item-count/2)*) |

The recommended strategy for annotating the factors and their levels using HED (as illustrated in Table 2) is to first create, for each level, a convenient HED “event concept” definition that includes a *Condition-variable* tag whose value is the factor name. The name of the definition is interpreted programmatically as the variable level for that factor (e.g., *Definition/**Famous-face-cond*** is a level for condition variable ***Face-type***). These elements appear in boldface in Table 2 to emphasize their role in documenting the experiment design. Notice that the BIDS event file excerpt in Fig. 2 (Section 1) includes a face_type column whose values (such as famous_face) give the factor levels.

The event file excerpt in Fig. 2 also includes a rep_lag column giving the number of trials past since the same image was first presented. This column includes numerical values only when the rep_status has values immediate_repeat or delayed_repeat, and n/a otherwise. Note that these values could be computed from the event table itself, but are included here (and in the accompanying W-H dataset submitted to OpenNeuro) to make that computation unnecessary.

Column-wise encoding of event (and experiment) design variables makes manual or automated extraction of the event design matrix from BIDS task events files straightforward. Here, the choice of column encoding for the face type and repetition status factors makes sense because the factor levels change with each face image presentation. When a condition variable has the same value for most (or all) events in the recording, using the event marker (row) encoding method to mark condition changes may be more appropriate.

The W-H experiment used a between-participants response-key assignment variable to control for handedness bias. The key assignment factor (with levels left_sym_cond and right_sym_cond) encodes the assignment of which index finger key press indicates the participant’s decision that the presented face is more symmetric than average. In the left_sym_cond condition, participants press a key with the left-index finger to indicate they perceived more than average facial symmetry, and press a key with the right index finger to indicate less than average facial symmetry. The left-right key assignment is counterbalanced across participants. Table 3 shows how to encode this key assignment using experiment control events.

**Table 3.** Encoding of the key-assignment condition variable using experiment control events (rows rather than columns of the task events files).

| Encoding in the task events file | Condition-variable level HED definitions |
| --- | --- |
| **Event file column:** `event_type`<br>**Column value:** `setup_left_sym`<br>**Annotation:** (*Def/Left-sym-cond, Onset*) | **Description:** Left finger key press signifies the presented face has above average symmetry (the right press, the opposite)<br>**HED:** (*Definition/Left-sym-cond, (Condition-variable/Key-assignment, ((Mouse-button, (Left-side-of, Computer-mouse)), (Behavioral-evidence, Symmetrical)), ((Mouse-button, (Right-side-of, Computer-Mouse), (Behavioral-evidence, Asymmetrical))))*) |
| **Event file column:** `event_type`<br>**Column value:** `setup_right_sym`<br>**Annotation:** (*Def/Right-sym-cond, Onset*) | **Description:** Right finger key press signifies the presented face has above average symmetry (the left press, the opposite).<br>**HED:** (*Definition/Right-sym-cond, (Condition-variable/Key-assignment, ((Mouse-button, (Right-side-of, Computer-mouse)), (Behavioral-evidence, Symmetrical)), ((Mouse-button, (Left-side-of, Computer-Mouse), (Behavioral-evidence, Asymmetrical))))*) |

Notice that key_assignment does not correspond to a column in the table of Fig. 2 (Section1). Because the level of this variable is constant for the entire recording, this variable is better encoded by inserting an *experiment control event* at the beginning of the recording to mark the *Onset* of this control-condition assignment. Here we insert an initial experiment control event with an event_type value of either setup_left_sym or setup_right_sym to encode the initial recording setup and key assignment. The onset time of this experiment control event is that of the first data point of the recording (see first event of the table in Fig. 2, Section 1).

Section 3 discusses in more detail how the definitions in Tables 2 and 3 can be used in conjunction with BIDS `…events.tsv` task events files to fully document the experimental design *within* the BIDS dataset annotation. HED tools now under development will then be able to automatically extract the design matrix and other statistics about the experimental design from HED definitions that include the *Condition-variable* tag and from experiment control events associated with these definitions.

## 3 HED annotation of a BIDS-formatted dataset

BIDS recommendations for archival data storage have quickly become a *de facto* standard for sharing raw neuroimaging data. This section demonstrates how HED event annotations are actually mapped into machine-actionable annotation of datasets organized according to BIDS specifications. A BIDS dataset typically holds data from an experimental study that includes a number of brain imaging data files recorded from one or more participants in one or more sessions and/or task or other conditions. BIDS specifies a particular dataset directory structure, file naming conventions, and permitted image data formats, making it easier for users and tool developers to access data without manual or computerized recoding.

In BIDS-formatted datasets, much of the metadata is located in `.json` (JavaScript Object Notation) text files called sidecars. File naming and folder architecture conventions associate the sidecar metadata with the data files. When the same metadata applies to many data files, BIDS allows metadata files to be placed higher in the dataset directory hierarchy. The metadata information is then inherited by data files in dataset sub-directories (the *BIDS Inheritance Principle*), thereby avoiding the need to repeat the same metadata within multiple files in lower levels of the BIDS folder hierarchy. HED leverages the inheritance principle by placing HED annotations in a JSON sidecar ideally at the top level in the dataset. HED tools are available to take concept tables such as those of Table 1 and Table 2 to automatically create a BIDS JSON sidecar for events files.

Table 4 below summarizes different mechanisms for including HED annotations in a BIDS dataset. The current case study includes HED information ONLY in the top-level …events.json sidecar file (shaded background) contained in the dataset root directory. That information is keyed to the column names of the individual `…events.tsv` files (Fig. 2 and Table 5 below) located at the lowest level of the dataset, each containing the list of event markers in the corresponding recording.

**Table 4.**
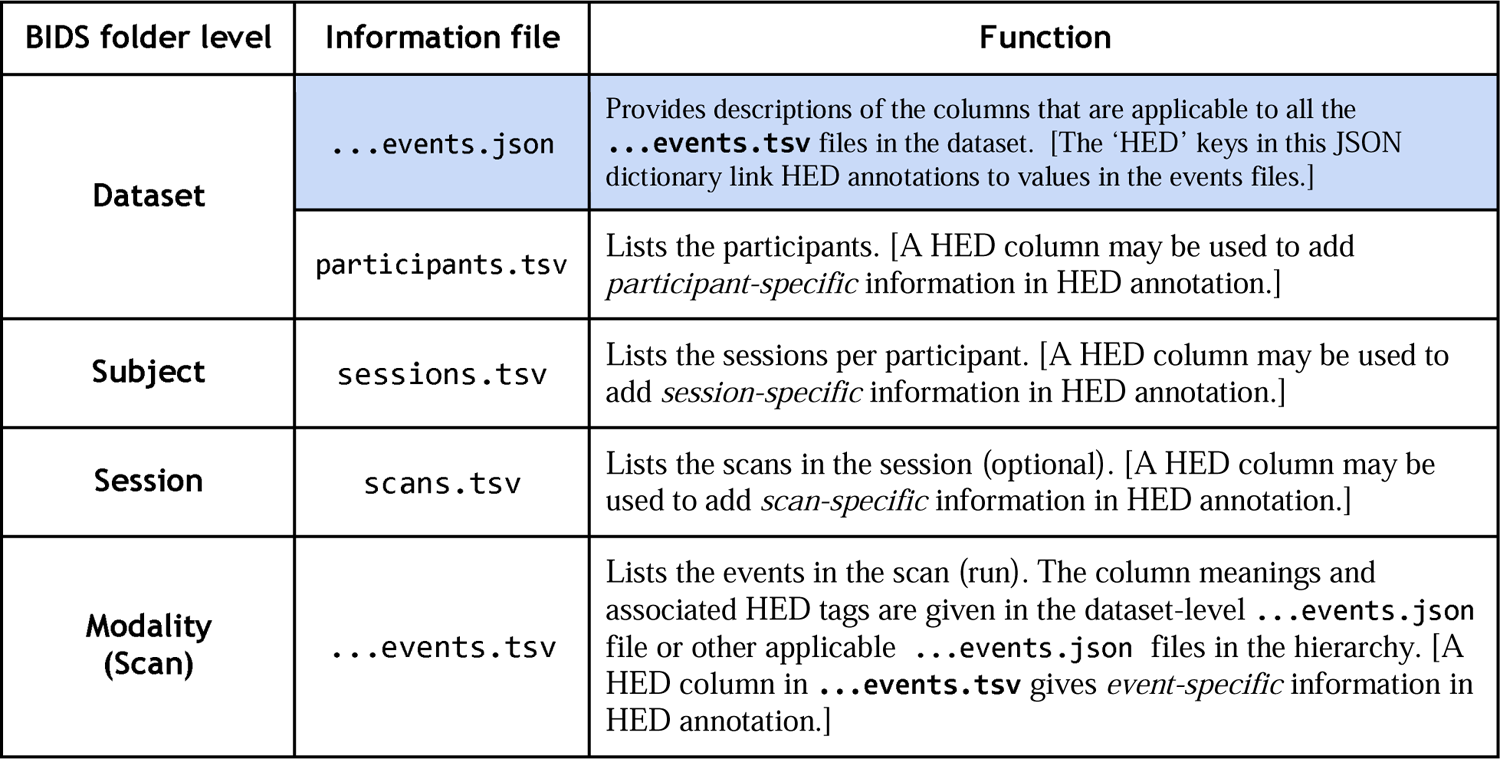
Mechanisms HED annotations in BIDS `.json` and `.tsv` metadata files. Many datasets may need only one `…events.json` file (blue background) placed in the top (Dataset) level folder.

| BIDS folder level | Information file | Function |
| --- | --- | --- |
| **Dataset** | `...events.json` | Provides descriptions of the columns that are applicable to all the `...events.tsv` files in the dataset. [The ‘HED’ keys in this JSON dictionary link HED annotations to values in the events files.] |
|  | `participants.tsv` | Lists the participants. [A HED column may be used to add *participant-specific* information in HED annotation.] |
| **Subject** | `sessions.tsv` | Lists the sessions per participant. [A HED column may be used to add *session-specific* information in HED annotation.] |
| **Session** | `scans.tsv` | Lists the scans in the session (optional). [A HED column may be used to add *scan-specific* information in HED annotation.] |
| **Modality (Scan)** | `...events.tsv` | Lists the events in the scan (run). The column meanings and associated HED tags are given in the dataset-level `...events.json` file or other applicable `...events.json` files in the hierarchy. [A HED column in `...events.tsv` gives *event-specific* information in HED annotation.] |

As summarized in Table 4, it is also possible to incorporate HED annotations in other BIDS `.tsv` files by including an extra column titled HED. These annotations are particular to the row of the file and should only contain HED strings (not HED definitions). For example, a HED string appearing in the HED column of `participants.tsv` pertains to the participant described in that row. In annotating more complex experiment designs, some HED information could be placed most efficiently in any or all of the four BIDS `.tsv` file types listed in Table 4 (if present) as well as in additional `…events.json` sidecars placed at lower levels in the dataset hierarchy, possibilities that for simplicity we do not discuss further here.

It is also possible to annotate individual events, or parameters that vary across individual events by recording additional individual-event HED tags in a HED column in the events files. Because of the difficulty in reading and editing annotations spread across individual events, this type of annotation should be avoided unless needed. However when, for example, presented stimuli have randomly varied properties (screen location, pitch, size, etc.), these details can be documented in this manner. Separate value columns in the event file with HED value annotations in the pertinent JSON sidecar can also be used to encode this information.

### 3.1 BIDS events.tsv files

At the lowest, single scan (data recording or run) level of the dataset folder hierarchy, BIDS event files are tab-separated value (`.tsv`) formatted text files with file names ending in `…events.tsv`. The BIDS naming convention associates the column headings in the `…events.tsv` event files with annotations contained in the relevant …events.json sidecar files – always including the top (full dataset-level) `…events.json` file. Here we use … prefixes in the filenames as placeholders for information embedded in the filename prefixes concerning data modality, task, session, subject, and run. The first line in a BIDS event file is a header line identifying each column, and each subsequent line corresponds to an event marker (an identified time point of interest within an identified event process) in the data.

Table 5 shows the excerpt of the BIDS event file of Fig. 2, color-coded to indicate the source of the expanded event annotations as shown in Table 7 (following, see Section 3.3).

**Table 5.** An excerpted BIDS `…events.tsv` file from the dataset displayed schematically in Fig. 2. The table includes the initial setup events as well as those defined in Table 1. Color-coded columns have relevant HED annotations defined in the `…events.json` sidecar file. Table 7 uses the same color-coding to dissect the expanded HED annotation of one of these events (the emboldened row in task events table below).

| onset | duration | sample | event_type | face_type | rep_status | rep_lag | value | stim_file |
| --- | --- | --- | --- | --- | --- | --- | --- | --- |
| 0.400 | n/a | 1 | setup_left_sym | n/a | n/a | n/a | 2 | n/a |
| 23.870 | n/a | 26275 | show_face_initial | famous_face | first_show | n/a | 7 | f032.bmp |
| 24.081 | n/a | 26488 | left_press | n/a | n/a | n/a | 256 | n/a |
| 24.750 | n/a | 27225 | show_circle | n/a | n/a | n/a | 0 | circle.bmp |
| 26.457 | n/a | 29095 | show_cross | n/a | n/a | n/a | 1 | cross.bmp |
| 26.940 | n/a | **29634** | **show_face** | **famous_face** | **immediate_repeat** | **1** | **8** | **f032.bmp** |
| 27.913 | n/a | 30701 | show_circle | n/a | n/a | n/a | 0 | circle.bmp |
| 27.990 | n/a | 30789 | right_press | n/a | n/a | n/a | 4096 | n/a |

**Table 6.**
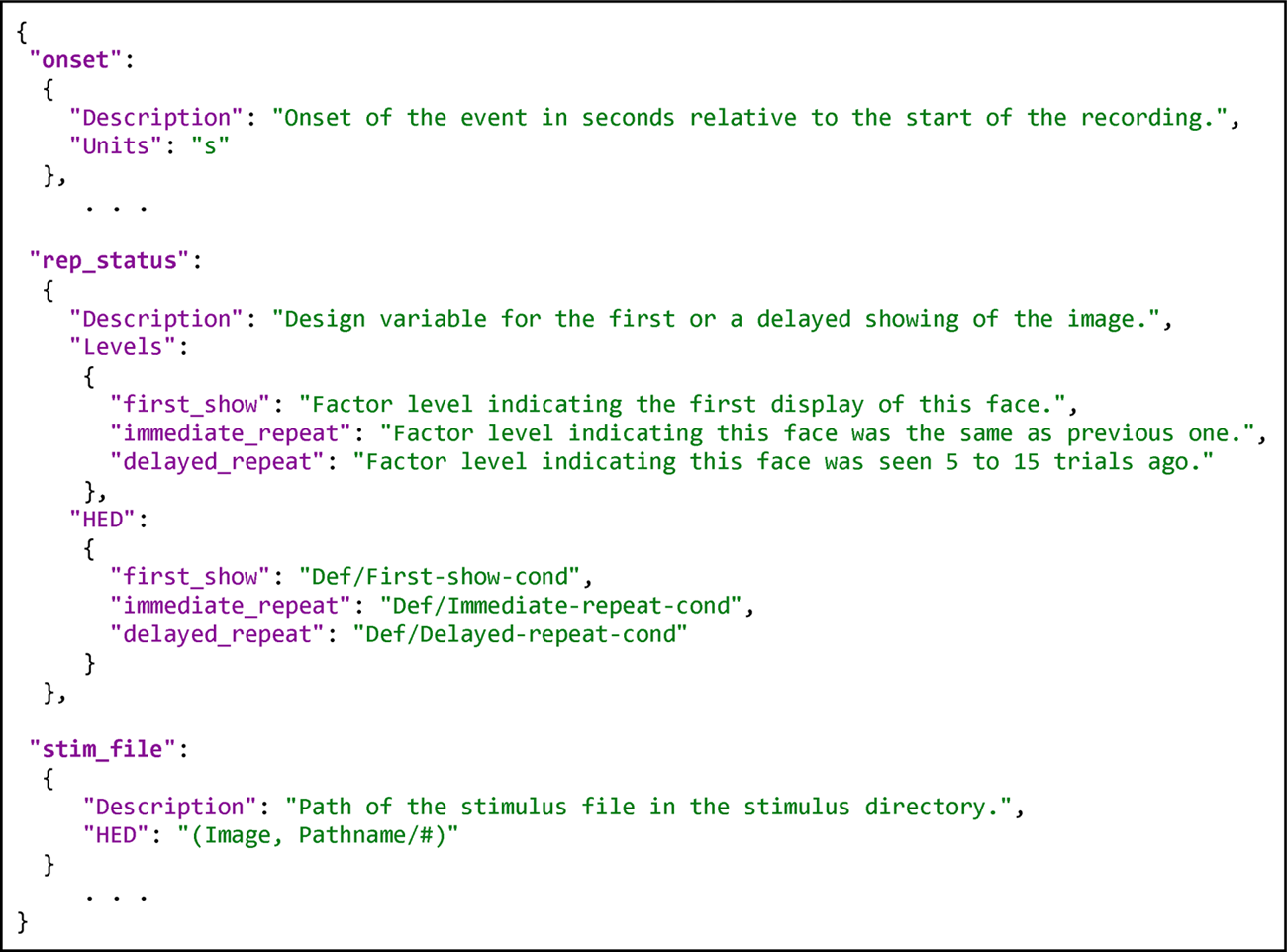
Excerpt of the top (dataset) level JSON sidecar file (`.events.jso`) for the W-H data.

**Table 7.**
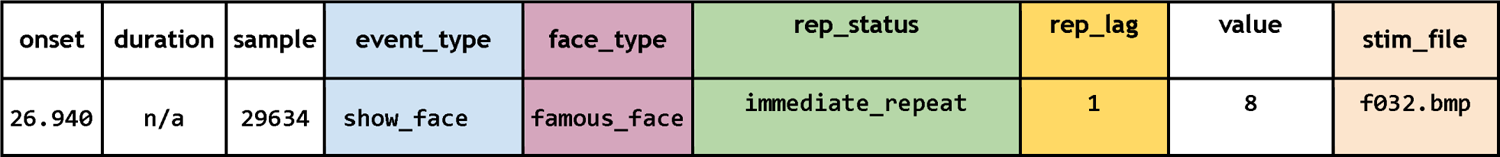
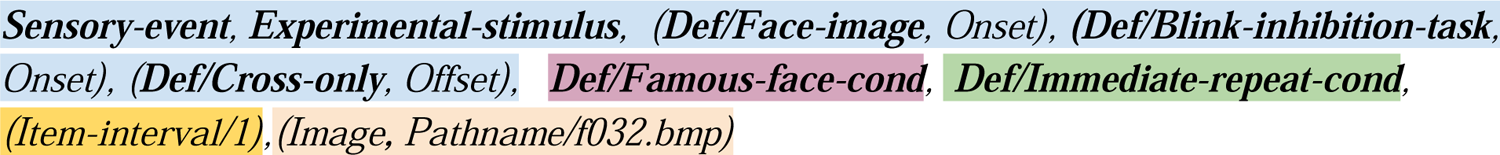
Assembled HED string for an immediate repeat of an image of a famous face (the seventh event in Table 5). The annotation also marks the end of the cross-only presentation and the onset of a blink inhibition period. The color coding of Table 5 is used to show the correspondence between annotation from the JSON sidecar file and the `…events.tsv` file column (event_type: blue, face_type: plum, rep_status: green, rep_lag: mustard, and stim_file: tan).

Note that Table 5 differs slightly from the events listed in Fig. 2 in that the second event has an event_type called `show_face_initial` rather than the show_cross of Fig. 2. As is often the case, the startup event in a block of trials differs from the internal block trials. The first reported event in all W-H recordings corresponded to the first showing of a face image rather than the showing of a fixation cross, although ERP analysis of the data suggests that this event actually occurred. Thus, the HED tags for show_face_initial includes *(Def/**Fixation-task**, Onset)* and does not include *(Def/**Cross-only**, Offset)*.

Each row in the task events file table gives information about a single event, typically functioning as a marker of the onset of an event process. BIDS requires event files to have onset and duration columns giving the onset time (in the data) and duration of each event in seconds. Users may add additional columns as needed. All columns in the task events file should be documented in one or more accompanying JSON-format sidecar files as described in the next section.

BIDS event files have two types of columns: categorical and value. **Categorical columns** allow a small number of distinct defined levels or categories, represented as text or numeric values. Other columns are **value columns**. Each row in the `…events.tsv` file in Table 5 has three categorical columns: `event_type` (blue), `face_type` (plum), `rep_status` (green) each with a relatively small number of distinct levels that will be annotated individually. Value columns in this file include `onset duration, sample, value` (all in white), and `rep_lag` (in mustard). The final `stim_file` column (tan) could be treated either as a categorical or as a value column depending on the number of distinct stimulus images. Here we treat `stim_file` as a value column because of the relatively large number of different face stimulus images used in the W-H experiment.

The distinction between categorical and value columns is important mainly because HED annotations are encoded differently for the two types of columns, as explained below. The column labeled value in the above example corresponds to the trigger values from the experimental control program and is retained for informational purposes. The columns displayed in white in Table 5 will not be annotated with HED.

### 3.2 BIDS events.json sidecar files

Many experiments can use a common and relatively simple event-design strategy that requires building only a single `…events.json` annotation file at the top level directory of the dataset to provide complete machine-actionable event annotation across participants and recordings when combined with the values in the individual recording `…events.tsv` files. In general, an organization using a single dataset-level `…events.json` sidecar is easier to annotate, understand, and maintain, so that is the organization we focus on here. The W-H annotation case study (Section 2) assumes that all the annotation of dataset events is in a single `…events.json` sidecar file (`task-FacePerception_events.json`) located in the top level tas dataset directory. Table 6 shows a portion of this sidecar file. See Supplementary Table 1 for the complete version.

For the complete version see Supplementary Table 1. The `…events.json` sidecar files are structured as dictionaries. The excerpt shown in Table 6 has three top-level keys (`onset, rep_status`, and stim_file) corresponding to column names in the `…events.tsv` file excerpt shown in Table 5. (Here the annotations for the columns sample, `event_type`, `face_type`, and `rep_lag` are omitted for readability but are included in Supplementary Table 1.) HED tools associate column metadata with particular columns in the event file using these column names. BIDS users may use additional top-level keys to include additional metadata in the JSON sidecars (e.g., the Levels and Description under `rep_status` in Table 6). We also use additional top-level keys to separate out the HED definitions for readability, although definitions may be included in the other annotations.

In Table 6, the metadata dictionaries associated with `rep_status` and `stim_file` have HED keys and hence include HED annotations. In contrast, the metadata dictionary associated with top-level key onset does not include a HED key, so it is considered to be an unannotated column and is ignored by the HED tools. If the HED key references a dictionary (as does `rep_status` in Table 6), HED assumes the task events table column is categorical, while if the HED key references a string (like stim_file in Table 6), HED assumes it is a value column. In either case, HED uses the corresponding HED key values to annotate the event.

Categorical column annotations in `…events.json` sidecar files include a separate HED annotation for each categorical value that appears in the corresponding column of the `…events.tsv` file (e.g., the categorical value first_show appearing in column `rep_status` of Table 5). Value column annotations (such as the one appearing for the stim_file column use a single HED string with a hash symbol (#) value placeholder to annotate the column. When the complete annotation for an event is assembled, the HED assembler tool replaces the hash symbol with the value from the respective row and column of `…events.tsv` file.

The next section explains how the annotation for an event is assembled by combining event information in the `…events.tsv` files with the HED annotations in the `…events.json` sidecar dictionaries.

### 3.3 Assembling and using the complete event annotation

HED assembler tools gather the BIDS `…events.json` sidecars applicable to an `…events.tsv` file and assemble a single HED string representing the annotation for each event marker (as represented by a line in the BIDS event file). The assembled HED string annotation for the second face display event ( w_face) in Table 5 is shown in Table 7. Parts of the HED string are color-coded to indicate which column annotation that portion corresponds to. The corresponding columns in the `…events.tsv` file of Table 5 use the same color shadings.

To annotate this show_face event (from the `…events.tsv` file excerpt of Tables 1 and 5), the HED assembler looks up the column annotations defined in the accompanying `…events.json` sidecar. As the onset, duration, sample, and value columns of the `…events.tsv` file do not have HED annotations in the `…events.json` sidecar file in this example, they are skipped. (Note: these columns could have been annotated as value columns). The show_face value in column `event_type` is translated into its HED definition (Table 1), then concatenated to the assembled annotation (light blue shading). Next, the annotation for famous_face in the `face_type` column is found in the sidecar and appended (plum shading). Then the category immediate_repeat in the `rep_status` column is looked up, and the corresponding HED annotation is included (green shading). Finally, the repetition lag value in the rep_lag column and the filename value in the stim_file column are substituted for the respective #’s in the corresponding annotations (mustard and tan shadings). The other column values are skipped in this process, because they have no HED keys in the `…events.json` sidecar dictionary.

During analysis, the HED tools can expand the definitions so that their values are available for searching and filtering. Supplementary Table 2 shows the assembled annotation of Table 7 in several forms, and demonstrates how the *Def-expand* tag is used with the substituted definitions to accomplish this expansion.

Combining the information in the BIDS `…events.tsv` files with the appropriate `…events.json` sidecar annotation file(s) enables powerful automated tools to be implemented. Given this information, such HED tools could automatically extract and optionally visualize the experiment task list, the underlying experimental design, and the temporal structure of a recording. Extensive statistics about the number of event markers with different properties could also be computed. Data could be separated into event-locked epochs with similar HED tags fitting a simple or complex description, and automatically bootstrapped to look for differences associated with different experimental parameters. Complex searches could be conducted across datasets (including datasets using different tasks and experimental designs) without need for manual re-coding. The case study developed in Section 2 and 3 illustrates the annotation process. The next section extracts “lessons to be learned” from this case study to formulate a set of “best practices” for event design and annotation.

## 4. Best practices in event design and annotation

A myriad of events, overt or covert, planned or unplanned, may unfold during the execution of an experiment. **How a researcher chooses to organize, report, and annotate events can completely change the capacity of a given dataset to support analysis, reuse, and reproducibility.** It may not be possible to record markers for every conceivable recordable event, nor may it be feasible to describe precisely their every detail. Incorporating fine details of all known events might indeed prove valuable to future analyses and mega-analyses. However, some limit in time and energy available must be accepted. One important strategy is to be sure to include the actual stimuli and/or virtual environments with the stored/shared data, as included here in the W-H data. Others wanting to exploit the analysis value of more detailed annotations of the data could then be in a position to add further details to the annotations. For example, the StudyForrest project (*studyforrest.org*) organized a team to more fully annotate events in the movie *Forrest Gump* that had been shown to participants in several neuroimaging studies.

***Event design*** as used here refers to the process of identifying, organizing, reporting, and sufficiently annotating the nature of events to a degree allowing complete interpretation of the event-related dynamics recorded during the experiment. The process includes listing the recurring types of event markers in the data, giving them easily recalled terms, and then defining each term using HED annotation. Ideally, these event markers and descriptions should include all that is relevant to both current, planned and future potentially fruitful analyses. *Event design should be the first step in augmenting a dataset with HED annotation*.

Best practice in event design encourages researchers to look beyond the immediate use of their data to broader questions. In particular: ***Which aspects are potentially important to future analysis*** (performed either by the data authors or others)? These analyses are likely to include meta-analyses and mega-analyses (Costafreda, 2012) (Boedhoe et al., 2019) (Bigdely-Shamlo et al., 2019) across shared datasets that may involve different designs, participant tasks, experimental conditions, and event types.

The event design process has two steps: first *identifying* which events to report or mark and then *mapping* the resulting event markers into usable annotations. **The most critical part of this process is recording and marking the events**, as events not marked in the data may not be recoverable. **Ideally, the event design process should be performed before data collection begins**, as the event design process clarifies what is being measured and whether those measurements can be used to achieve experimental goals. In any case, most of the information required by a good event design process will be required in publications reporting the work, so performing a preliminary event design can help to assure that important details are not confused or overlooked later. In this section, we discuss the event design process and suggest guidelines for it using the W-H dataset as a case study. Even when HED annotation is performed after data collection, beginning the annotation process with event design is useful for deciding how to best annotate the data.

### 4.1 Event design for the W-H experiment

The W-H event design developed in Sections 2 and 3 above is not the one distributed with the original shared OpenNeuro dataset *ds000117*, but was developed based on the recommended event design practices with the generous assistance of the data authors Wakeman and Henson to make additional event type and timing information available in the data. The MEEG data of the redesigned dataset are available as OpenNeuro dataset *ds003645*. The event design of Table 5 marks the onsets and offsets of all the experimental stimulus sensory presentations and participant action motor responses using the annotations and encoding of the `event_type` column of Table 1. Further, the 3 × 3 experimental design is represented (using information in the `face_type` and `rep_status` columns and the encoding described in Table 2.)

Table 5 defines a setup_left_sym experiment control *meta-event* whose time is that of the first data sample. This meta_event can also be used to store other annotations applicable to the entire recording, such as the visual presentation screen size and participant distance (as available) (Table 3). Since the (left = ‘symmetric’) key assignment is in effect for this entire recording, it is more efficient and clearer for tools to encode it as an initial meta-event rather than giving it its own column in the `…events.tsv` files requiring the same value to be repeated for every motor response event. If we want to use the single JSON events sidecar at the top level in the BIDS dataset file hierarchy, every value in the `…events.tsv` files must have the same meaning across the entire BIDS dataset. A setup_right_sym meta-event must also be introduced there to apply in the datasets using the (right = ‘symmetric’) key assignment.

The event table also includes a column labeled sample that gives the data sample number of the event marker. This column is recommended in the BIDS standard and is good practice since the precision of the onset values is left completely open in BIDS and accurate event timing is extremely important for MEEG analysis. The value column is here not necessary, because its information is already encoded in the `face_type`, `rep_status`, and rep_lag columns, but we have retained it to maintain the connection with the original shared dataset, since the value column captures the actual event code triggers produced by the experiment control software.

For comparison, Table 8 shows a sample of the event file for the MEEG portion of the W-H data, as originally shared. The `…events.tsv` files only give the onsets of the face presentations and contain no markers for other sensory presentation or participant responses, limiting the usability of the data for analysis, further analysis, and meta/mega-analysis.

**Table 8.** MEEG event file for run 1 of session 1 of subject 01, as originally shared.

| onset | duration | onset_sample | stim_type | trigger | stim_file |
| --- | --- | --- | --- | --- | --- |
| 24.2073 | 0 | 26628 | Unfamiliar | 13 | meg/u032.bmp |
| 27.2473 | 0 | 29972 | Unfamiliar | 14 | meg/u032.bmp |
| 30.3545 | 0 | 33390 | Unfamiliar | 13 | meg/u088.bmp |
| 33.3618 | 0 | 36698 | Unfamiliar | 13 | meg/u084.bmp |

Table 8 is considerably shorter and narrower than Table 5 (our recommended version), but it is missing critical information (e.g., `rep_status` and all the events marking presentations of the fixation cross and focusing circle, as well as the key press events). Difficulties introduced for downstream analysis by not recording and reporting all possible sensory and participant action events are discussed in more detail in Section 4.3 and Section 4.4, respectively.

Another difficulty in Table 8 is the use of non-orthogonal encoding of the experimental design in the event-recording hardware system trigger column, whose 12 distinct values are shown in Table 9.

**Table 9.**
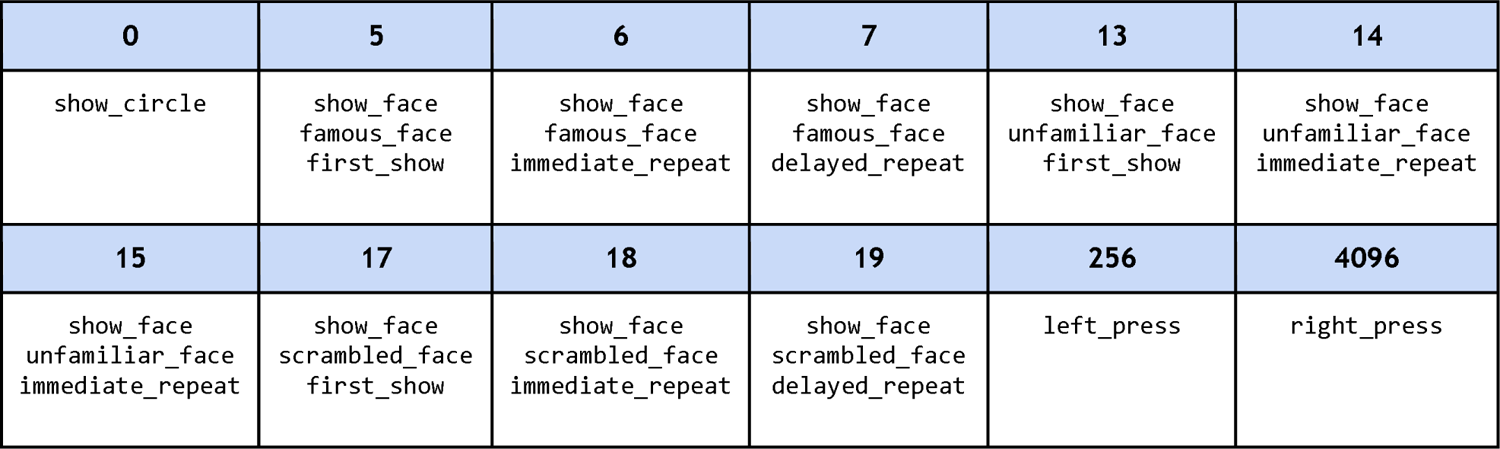
The 12 trigger values from the original W-H (shaded rows) and their respective interpretations.

| 0 | 5 | 6 | 7 | 13 | 14 |
| --- | --- | --- | --- | --- | --- |
| show_circle | show_face<br>famous_face<br>first_show | show_face<br>famous_face<br>immediate_repeat | show_face<br>famous_face<br>delayed_repeat | show_face<br>unfamiliar_face<br>first_show | show_face<br>unfamiliar_face<br>immediate_repeat |
| 15 | 17 | 18 | 19 | 256 | 4096 |
| show_face<br>unfamiliar_face<br>immediate_repeat | show_face<br>scrambled_face<br>first_show | show_face<br>scrambled_face<br>immediate_repeat | show_face<br>scrambled_face<br>delayed_repeat | left_press | right_press |

While it is possible to tag each trigger value as in Table 9 to associate it with the factors and levels it represents, the non-orthogonal or mixed encoding used to build the trigger codes makes downstream analysis much more likely to require manual re-coding, thereby making the dataset difficult to include in further analysis. In the recommended design (Table 5) the independent factors `face_type` and the `rep_status` are represented by independent columns in the events file, making it easy for automated processing to detect the 3 × 3 design. Encoding of experimental conditions is discussed in more detail in Section 4.5.

### 4.2 Pitfalls in reporting events by-trial rather than by-event

An overall guideline for reporting events strongly favors expressing each relevant event with its own (onset) event marker and corresponding line in the event file. Where relevant, offset time information for events representing processes with appreciable duration should also be reported. In some cases, event markers for intermediate points of interest in an event process may also be important for analysis, for example onsets of individual syllables in spoken words or critical points of hand/arm movements in reach trajectories. HED also supports use of such markers, though we have not here given an example of their use.

##### Guideline 1: Event files should be organized by event.

Event files should report one event marker per line. Event files should contain markers (lines) for all onsets and offsets of relevant sensory stimuli, motor actions, participant tasks and task conditions, condition changes during the recording, time organization, plus setup meta-event information organized during event design. When computation of response times or delays, or results of other computations on the basic event data are stored in a column added to an event table, the event table should still include rows representing the onsets and offsets of the actual framing events used to compute these response times or delays.

While this recommended *by-event* organization may seem logical, currently many shared BIDS datasets instead use a *by-trial* organization or some hybrid organization. *By-trial* organization treats each *trial* as a single event that is given one row in the event file, and expresses all other relevant trial event markers in that row as offsets from the trial latency in the data. Such *by-trial* organization has many disadvantages for event-related and more general analysis approaches, most prominently a lack of clarity with respect to the timing of other MEEG data-influencing events. As an illustration consider the sample of an event file originally shared for the fMRI portion of the W-H experiment shown in Table 10.

**Table 10.**
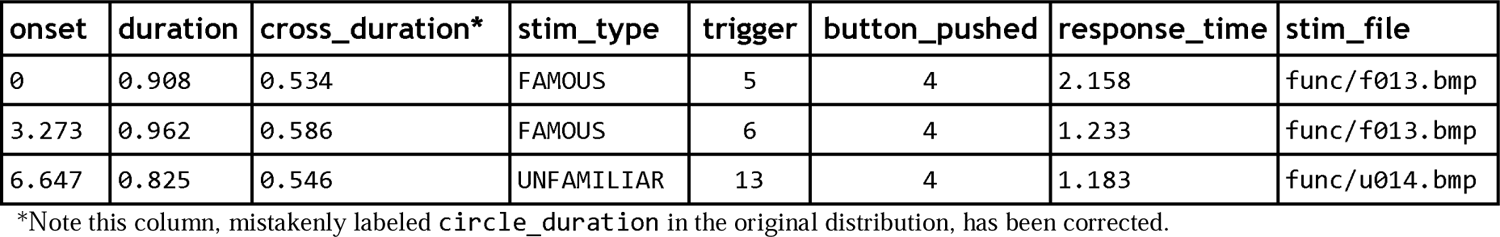
The W-H experiment fMRI event file for the first run of session 1 for subject 01, as originally shared.

| onset | duration | cross_duration* | stim_type | trigger | button_pushed | response_time | stim_file |
| --- | --- | --- | --- | --- | --- | --- | --- |
| 0 | 0.908 | 0.534 | FAMOUS | 5 | 4 | 2.158 | func/f013.bmp |
| 3.273 | 0.962 | 0.586 | FAMOUS | 6 | 4 | 1.233 | func/f013.bmp |
| 6.647 | 0.825 | 0.546 | UNFAMILIAR | 13 | 4 | 1.183 | func/u014.bmp |

\*Note this column, mistakenly labeled `circle_duration` in the original distribution, has been corrected.

When motor response events are reported only as response_time delays, it is not always clear whether the time is relative to the trial anchor event or to some other event. Events that occur before the anchor event are not always expressed with a negative delay (e.g., here cross_duration is positive, although the cross display occurs *before* the anchor face presentation onset event). While it is possible to calculate the onsets and offsets of the visual stimuli from the various durations and response times relative to the anchor event, a data user would have to do a very careful analysis of the documentation and published papers to correctly identify the sensory and motor action event onsets and offsets. Performing this anew for each shared dataset in any future mega-analysis across shared datasets would be infeasible -- or at best heroic.

By clearly identifying *all* experimental sensory events in a column named `event_type` or something similar, the design of Table 5 makes processing much easier. To reiterate, ***identifying all event onsets and offsets is increasingly important for many analyses, in particular those that use standard or new methods to model the complex, interacting effects of events on cognition and MEEG dynamics*.**

A second issue with *by-trial* organization of an event table is its lack of extensibility. For consistency, each row in *by-trial* reporting should contain information about the event sequence for the trial. Often however, conditions change and other events need to be recorded outside the strict *by-trial* structure, thereby complicating the annotation process. When later adding event markers (lines) to the event file to identify additional events in the data (such as blinks, alpha spindles, interictal spikes, background noise or outbreaks), researchers must decide whether to add additional columns and express the new times in terms of trial offsets or to add additional rows and treat the new markers as separate non-trial events. The difficulty with the latter approach is that the marked event times are likely to cross trial boundaries, thus requiring dataset-specific manual coding and analysis to unwind the information about those events. Operations such as regressing out the effects of overlapping events or determining effects of ongoing event context cannot be performed without first obtaining a distinct, well-ordered record of the dataset event onsets and offsets.

### 4.3 Documenting sensory presentations

##### Guideline 2: All known sensory presentations that are intended to or may affect neural responses should be marked and annotated.

Sensory presentations (including their onsets and offsets), as well as transitions between trial, performance blocks, stimulus or task condition changes, and other known or easily computed significant moments) should be given event markers. In addition to the formally designated experiment “stimuli,” dataset sensory presentations may include delivery of instructions, feedback, auxiliary stimuli including fixation points, cues, other filler images, changes in background, plus any unplanned events noted as having occurred during the recording. The role of each sensory presentation within the task and experiment, as well as a description of the sensory presentation and modality itself should be documented.

**Event annotation should aim to document all that the participant experiences.** At a minimum, thoughtfully detailed reporting of participant sensory experience allows analysts to regress out the influences of other sensory presentations on dynamics associated with presentations of the primary stimuli; nonlinear modes of analysis may benefit still more from this information, quite possibly in ways yet undocumented.

As first shared, the shared W-H MEEG dataset noted only face image presentation onsets, while the fMRI dataset also included cross duration and key press response times as well as indicating which key was pressed (left or right). Papers published by the authors on the fMRI dataset also included a somewhat more complete description of the event sequence depicted by the timeline of Fig. 2.

We found some ambiguity in the published description of the W-H MEEG experiments. When did the first trial begin? Did recording begin at the start of the first trial? If not, was a white circle displayed at the beginning of the recording? To avoid such ambiguities, it is best practice to **write experiment control scripts that automatically output event markers for every sensory presentation event** as well as the data itself.

### 4.4 Documenting participant responses

##### Guideline 3: Participant motor responses (and any other recorded participant actions) should be reported.

Instructed participant responses or actions should be marked as individual events (or event sequences) rather than reported only as reaction times and/or by noting the category of the participant response (e.g., for the W-H experiment, only noting responses as having indicated a ‘symmetric’ or ‘asymmetric’ judgment). Motor actions themselves, their planning, and accompanying and ensuing assessment processes are all supported by brain dynamics that are very likely to be reflected (in part) in neuroimaging data features.

*As with sensory presentations, motor responses should be first annotated from the perspective of **what the participant does**, not what it means in terms of the experiment design and task. At a minimum, the annotation should document who acts and what action they take. The experiment control program’s handling of correct, incorrect, and omitted response actions (if computed by it) should also be articulated if these affect the selection of later stimuli*.

*Other types of participant actions, instructed or incidental, should also be documented using appropriate vocabulary from the HED base schema. If these actions were not instructed, they are not likely to be part of the initial experiment design, so they need to be entered as data features post hoc*.

In the W-H experiment, participants were instructed to press one of two keys with their respective left or right index fingers to indicate their assessment of the ‘symmetry’ of the presented faces. This symmetry evaluation task was unrelated to the experimenters’ own true objective in running the experiment. Perhaps for this reason, the participant responses were not fully documented in the W-H data as originally shared, and there was no indication in the dataset documentation of what would occur when or if the participant withheld a key press entirely.

Thinking more broadly about potential further uses for the data (e.g., when building the event design) may hopefully inspire data authors annotating their data to make it fit for a broader range of uses and sharing, thereby considering it worthwhile to add all available detail about subject performance to the shared dataset to enhance continued dataset usability. Here, for example, the W-H face symmetry evaluation task might itself be of some future interest, as might be how the pose or gender of a presented face affects brain dynamics and motor responses. Such readily recorded variables might also be treated as dependent variables to strengthen the statistical reliability of effects of interest in any analysis of the data.

### 4.5 Documenting experimental conditions, controls, and designs

##### Guideline 4: Experimental conditions, both fixed and changing, should be identified, whether they are part of the experimental design or are put in place to control experimental bias.

All experimental conditions should be documented, not just the main design variables. Full documentation allows researchers to systematically test for statistical differences in data features under various conditions. The explicitly stated experimental design provides the obvious factors to be annotated.

**Any aspect of the experiment that was controlled for bias can provide a condition for annotation.** Elements that are counterbalanced or randomized in a specified range should always be given serious consideration for explicit annotation as experimental conditions.

The span of each condition should also be identified. Was the condition varied by trial, by block, by run, by session, or by participant? If so, how and when – precisely?

In addition to the experimental conditions encoded in Tables 2 and 3, the W-H dataset has other potential condition variables such as the face image sex (with levels *female* and *male*), to encode the perceived sex of the presented faces. There is a large literature on the relevance of sex/gender in face recognition (Mishra et al., 2019), and the dataset description mentions that 50% of the stimulus faces were female and 50% male. The sex of the study participants was recorded; it would also be possible to identify, record, and annotate the sex of the faces in the shared stimulus images. One could then for example ask whether sex of the imaged face influenced judgment response time or any MEEG data feature.

### 4.6 Task specification

##### Guideline 5: All explicit as well as implicit participant tasks should be identified.

A participant task is an organized participant activity performed during (or sometimes before or after) the experiment that may influence participant brain dynamics. **Explicit tasks** usually (though not always) determine and lead to actions that the participant performs (or inhibits) intentionally during the experiment – and **should always** be documented. **Implicit task challenges**, whether or not directly reflected in participant actions, **should also** be documented – particularly if they are part of the experimental design. Explicit pre- or post-session tasks **external** to the recording session (often an aspect of experiments on learning or memory, for example) may also be considered for annotation, as in such experiment designs they may be intended to produce residual or priming effects in the session data.

#### Explicit tasks

The W-H experiment has three instructed or explicit tasks: face symmetry evaluation, gaze fixation, and blink-inhibition. The face symmetry evaluation task was the primary explicit task that the experimental participants were told to focus on. However, in the original data evaluation plan, this task was chosen solely to direct participant attention to each face and was irrelevant to the actual scientific goals of the experiment. Because this explicit task was the central activity the participant was instructed to perform, it should be documented as an explicit task (even if, as here, it did not enter into the original data evaluation plan).

As is common with many neuroimaging experiments, the W-H experiment instructions also included two other explicit tasks: blink inhibition and gaze fixation. Participants were asked not to blink when a face was being shown and were also told to fixate their gaze on the cross when visible.

Intentional fixation not only reduces the extent of natural eye movements but also may impose an additional mental load on participants. Instructed participant actions that may affect the recorded brain dynamics including, here, blink inhibition (Shultz et al., 2011) (Berman et al., 2012) and fixation (Stacchi et al., 2019), should always be considered explicit tasks for annotation. At a minimum, future analyses of the W-H dataset might test how successful participants were in inhibiting blinks during the specified period. Failures to inhibit might also be linked to variation in the recorded brain dynamics.

The separation of the two eye activity-related tasks into distinct tasks is necessary for the W-H dataset because the blink inhibition task applies only while the face image is being displayed, while the gaze fixation task is active during both the pre-stimulus interval and the face image presentation. Thus, these instructed intentions (affecting action) must be documented as separate tasks. While blink inhibition and gaze fixation could be annotated as experimental conditions in Table 2, activities performed intentionally by participants should be annotated as tasks, while elements that correspond to the setting and varying of experimental parameters should be annotated as experimental conditions or controls supporting interpretation of experiment control events and mega-analyses across datasets recorded under different conditions.

The W-H fMRI sessions also included data from a behavioral *face-memory test* conducted after the imaging session was completed. Since the participants did not have foreknowledge of the behavioral test, an experimental note to this effect should be included in the annotation of those data to inform further analysis. In the post-imaging *face-memory test*, W-H asked participants to view face images and to record whether they remembered seeing the face in the experiment sessions. These responses were not included in the original shared W-H dataset. To include them, BIDS conventions expect that they be stored as a third, behavior-only, W-H experiment session. This behavioral data is included in the new W-H-MEEG dataset available on OpenNeuro.

### Implicit tasks

The inclusion of repetition status as a design variable indicates that the experimenters were aware that detection of face novelty (or repetition) was very likely associated with brain dynamic effects in these data. The repetition status factor helps users assess the influence of this design factor in the data. The detection of face novelty might thus be considered to be an implicit task, that is, an activity that the participants were not directly instructed to perform, but rather could be expected to perform (either intentionally or near-automatically) during the course of the experiment, or at very least, that could affect the recorded brain dynamics in some systematic manner. The repetition status design variable could also be associated with another implicit task, face recall, as repeated-face recognition and new-face novelty detection are associated with distinct brain activity patterns (Debener et al., 2005) (Murashko & Shmukler, 2019) (Courchesne et al., 1975).

The `face_type` design variable, indicating whether the image is of a famous face, an unfamiliar face, or a scrambled face, is also an obvious candidate here for implicit task designation. The mixed presentation of these three rather different sets of images can be expected to have posed one or more implicit task demands on most or all of the participants. Here, possible implicit tasks include *nonface recognition*, *known face recognition*, *unknown face appraisal*, and *known face identification*. Here the scrambled face (*nonface*) images were a (⅓) minority of the presented stimuli and differed markedly from the other face stimuli in visual presentation. Neuroimaging responses to novel, outside-expected-category stimuli have distinct and long-known features.

Clearly, potentially a large number of implicit tasks could be annotated for analysis of these data. The choice of how to identify and annotate implicit tasks depends on what the annotator thinks may be of value to explore or test in the data. Very often, implicit tasks are associated with experimental control variables for experimental design or bias control. Even when an implicit task has no direct indication of whether the user actually performed the task, the annotation can be useful for directing downstream users of the data towards aspects of the experiment that are or may be associated with effects in the data or when comparing differences in effects across experiments.

By annotating such implicit tasks, shared datasets become amenable to future cross-dataset meta-analysis (of computed data features) and mega-analysis (of the raw data). We anticipate that common best practice norms will develop gradually as researchers see the value added to their data by performing the annotation in a style compatible with other shared datasets involving different experiment and task designs.

### 4.7 Documenting temporal organization and architecture

##### Guideline 6: The temporal architecture of each recording should be annotated.

The internal **temporal architecture of each recording should be documented**, including timing of performance blocks and rest periods between task blocks. If blocks of trials were used to vary or counterbalance some aspect of the experiment, event markers for the beginnings and ends of these blocks should also be included. More generally, **information that remained fixed throughout the recording should be gathered and annotated using a meta-event marker inserted at the time of the first data sample.**

Many neuroimaging datasets are organized into blocks of continuous or repeated task performance interspersed with rest periods. The W-H MEEG recording sessions were organized into 6 runs of 7.5 minutes duration containing between 140 and 150 face stimulus presentations (and thus, trial event sequences). Within each run, the W-H MEEG data do not have an explicit block structure beyond the trial level, though other experiments may have temporal structure within runs imposed to counter-balance various experimental factors.

A review of the W-H MEEG metadata showed that between 3 and 6 minutes elapsed between MEEG session runs. Analysts assume that electrode caps or other sensors were not repositioned between runs *in the same session*. If this was *not* the case, the information should be clearly marked in the data, typically by separating it into separate *data sessions* in which channel locations do not (or are assumed to not) vary. Head movements with respect to the MEG dewar and its embedded sensors are a key concern in MEG studies, and movement files acquired at 1- second intervals are available for the W-H MEEG dataset.

Although the W-H experiment does not have a particularly complex temporal architecture, the authors do use the concept of an experimental trial, so a definition *(Definition/Trial, (Experimental-trial))* could be included in the annotation to indicate the onset and offset of these trials, when this would seem useful for planned analyses. The distributed BIDS task event data includes a Trial column to make the grouping of the events in each trial more clear. Note however our cautions (in Section 4.2) about annotating events only in relation to trial event groupings.

### 4.8 The event design process

1. **Sketch a rough time-line (as in Fig. 2)**. Having a good picture in mind of how the experiment unfolds is a helpful starting point.
2. **List the basic event concepts of the experiment and give them concise, easily interpretable names.** Relevant concepts include sensory presentations, participant tasks and motor and/or verbal responses, experiment design, and bias control factors.
3. **Write a concise but complete text description of each event concept**. A good starting point is to create a table of component names and descriptions.
4. **List the needed event marker types** (as in Fig. 2), including Onsets and Offsets plus any other.
5. **Assign a primary HED *Event* category tag** to each marker (as in Fig. 3).
6. **Determine which additional columns if any** should be in the BIDS `...events.tsv` file.
7. **Verify that the event concepts (stimuli, responses, factors, levels, tasks)** can be associated either with `…events.tsv` event table markers (rows) or with event table columns having HED definitions in the `…events.json` files.
8. **Check and iterate** as needed.

Event design is usually an iterative process. Below are suggested steps to maximize the chances that the design leads to complete and valuable annotation:

In performing event design, annotators should initially not try too hard to complete detailed HED tags, but should make sure that the relation of the event markers to the experiment structure is correctly expressed. Detailed event annotation can be easily added (or edited as needed) later in the process by editing the `…events.json` files.

## 5. Discussion and roadmap

Good event design and annotation are essential for ensuring the usability and longevity of both shared and stored neuroimaging data. Researchers need to think beyond the immediate problem to be analyzed and think about how to share data in a manner that allows other researchers to rely on the data and benefit their research by its use. Many publishers encourage researchers to publish their data in a publication distinct from the primary published work. Separate publication increases the visibility of the work and provides authors with the opportunity to produce data with high quality documentation.

Current standards and conventions for sharing neuroimaging data including BIDS focus on file structure and inclusion of basic metadata but have few requirements with respect to annotation of experiment events. In fact, we know of no system other than HED that supports annotation of the detailed nature of *events* in human neuroimaging time series data. Many of the BIDS-validated MEEG datasets that we have evaluated on OpenNeuro have sparse or missing event annotations (Delorme et al., 2020). For such BIDS datasets, adding a single `…events.json` sidecar file, as illustrated here, or improving an existing one may be all that is needed to turn an otherwise impoverished and unusable dataset into a richly informative one.

Annotators should begin by simply naming and describing sensory presentations, participant response actions, explicit tasks, and task conditions. Even without including very detailed HED tags in the definitions of these concepts, their presence in the annotation can allow future automated tools to produce detailed informative dataset summaries and structural information. For example, the presence of *Condition-variable* tags allows tools to extract information about that condition variable even if no other tags are provided. Additional details can be added to the `…events.json` file at any time without modifying the rest of the dataset.

Ideally, a thoughtful approach to event design as defined here should be initiated before the experiment begins. The reported event streams should be unwound so that each event phase is reported (*by-event*) in its own row in an `…events.tsv` file rather than having some event phases being reported indirectly as offsets or response times relative to other reported events (Section 4.2). The latter (*by-trial*) approach can result in hopelessly convoluted event streams, particularly when additional data-feature or expert-annotation events are added *post hoc*. Such reporting makes analyses as simple as regressing out the effects of overlapping temporal events nearly impossible without extensive manual re-coding specific to each dataset.

### HED Library Schema

HED now supports library schema, specialized HED vocabulary trees used when needed for an annotation in conjunction with the HED base schema for annotation terms needed by specific research user communities and applications. Currently, a SCORE library schema for standard labeling of neurophysiological clinical EEG recordings (Beniczky et al., 2017) is under development, and work is beginning on a MOVIE library schema for annotating experiments involving 4-D (animated) stimulus presentation. A linguistics library schema is under consideration by another group. We are ready to assist any interested user groups in developing library schemas to make available specialized subfield annotation vocabularies available in HED, for example those needed to describe experiments involving biomechanics, virtual reality, music, or other research areas.

We also expect to make more progress on difficult remaining annotation issues including documenting spatial relationships, body movement frames, and task designs in HED. We also plan to work with experiment control program developers to investigate approaches for adding HED tags to experimental events and recorded participant actions during data acquisition. We look forward to documenting and demonstrating the value of the HED context framework, only briefly discussed here (Section 2.4), for performing context-aware analysis of neural dynamics.

HED tools for validation and analysis support, some already implemented and others now under development, are being written in Python. A HED JavaScript validation tool has been incorporated into the official BIDS validator and is being continually improved. Online tools are available at *hedtools.ucsd.edu*. The CTagger annotation tool (*github.com/hed-standard/CTagger*) provides a simple-to-use interface that supports “learning through doing” HED annotation. HED support tools for MATLAB have also been incorporated into EEGLAB including tools to select and process data epochs based on searches through dataset HED annotations. All HED code and issue forums are available on the HED organization GitHub website (*github.com/hed-standard*). The HED specification and list of tools and resources is available at *hed- specification.readthedocs.io/en/latest/index.html*. Further documentation is available on the HED website (www.hedtags.org).

Finally, we should not ignore the suitability for HED annotation to be applied equally well and in the same manner to events in other time series data including fMRI. The sensory presentations and participant actions, as well as in-data changes in experimental parameters and conditions in the many thousands of reported fMRI experiments are as equally well suited to HED annotation as are the (typically quite similar) experiment events in many MEEG experiments.

We believe that the time has now arrived for widespread recognition and acceptance of the need for a common framework for performing event annotation of neuroimaging time series data that facilitates replication as well as advanced analysis, either within or across experiments and datasets. Third-generation HED and its supporting tools are now in open release, (github.com/hed-standard). We welcome reader comments, suggestions, and participation.

## Acknowledgments

We would like to express our deep appreciation to Daniel Wakeman and Richard Henson for sharing their rich and useful dataset with the human neuroscience research community, for their patience in helping us to understand the experimental details, and for their willingness to dig out additional information for us to include and discuss here. We would also like to acknowledge the seminal vision and contributions of Nima Bigdely-Shamlo, who initially conceived the HED system concept and demonstrated its potential for practical use and importance for data search and analysis within and across studies (Bigdely-Shamlo et al., 2013). We also thank Jonathan Touryan of the Army Research Laboratory and Tony Johnson of DCS Corporation for their work in supporting the development of HED for EEG data sharing. Ian Callanan and Alexander Jones are primary tool developers of the HED-3G supporting infrastructure. This project received support from the Army Research Laboratory under Cooperative Agreement Number W911NF- 10-2-0022 (KR) and from NIH projects R01 EB023297-03, R01 NS047293-l4, and R24 MH120037-01 (SM). The Swartz Center for Computational Neuroscience is supported in part by a generous continuing gift from The Swartz Foundation (Old Field, NY).

## Author contributions

Conceptualization: KR, SM, DT Methodology: KR, SM, DT Writing – original draft: KR, SM, DT, SA, AD Writing – review & editing: KR, SM, DT, SA, AD Data curation: DT, KR, AD Software: KR, DT, AD, SA Visualization: SM, KR

## Ethics

All data used in this study are publicly available.

## Data availability

https://openneuro.org/ds003645.

---

**Supplementary Table 1:**
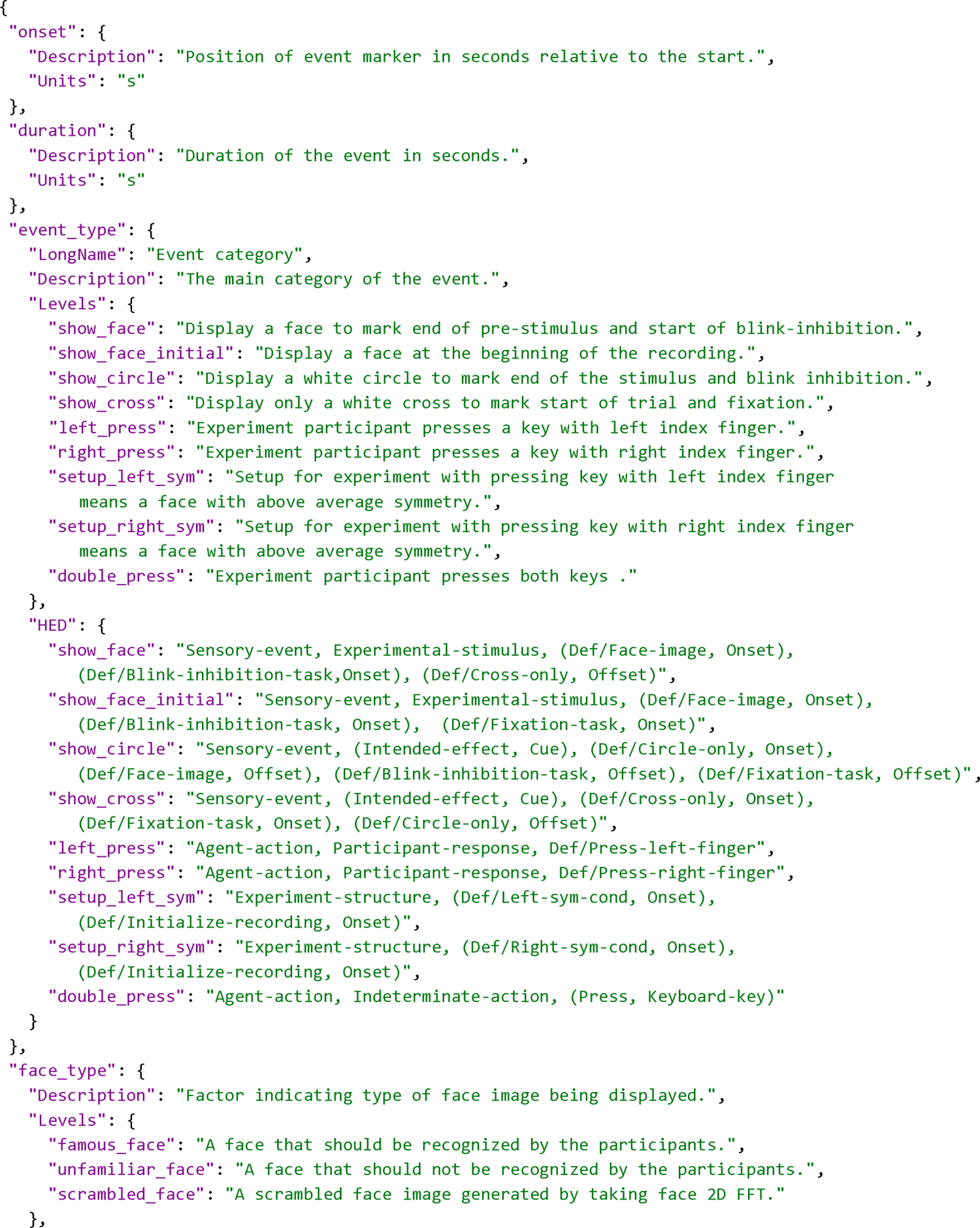
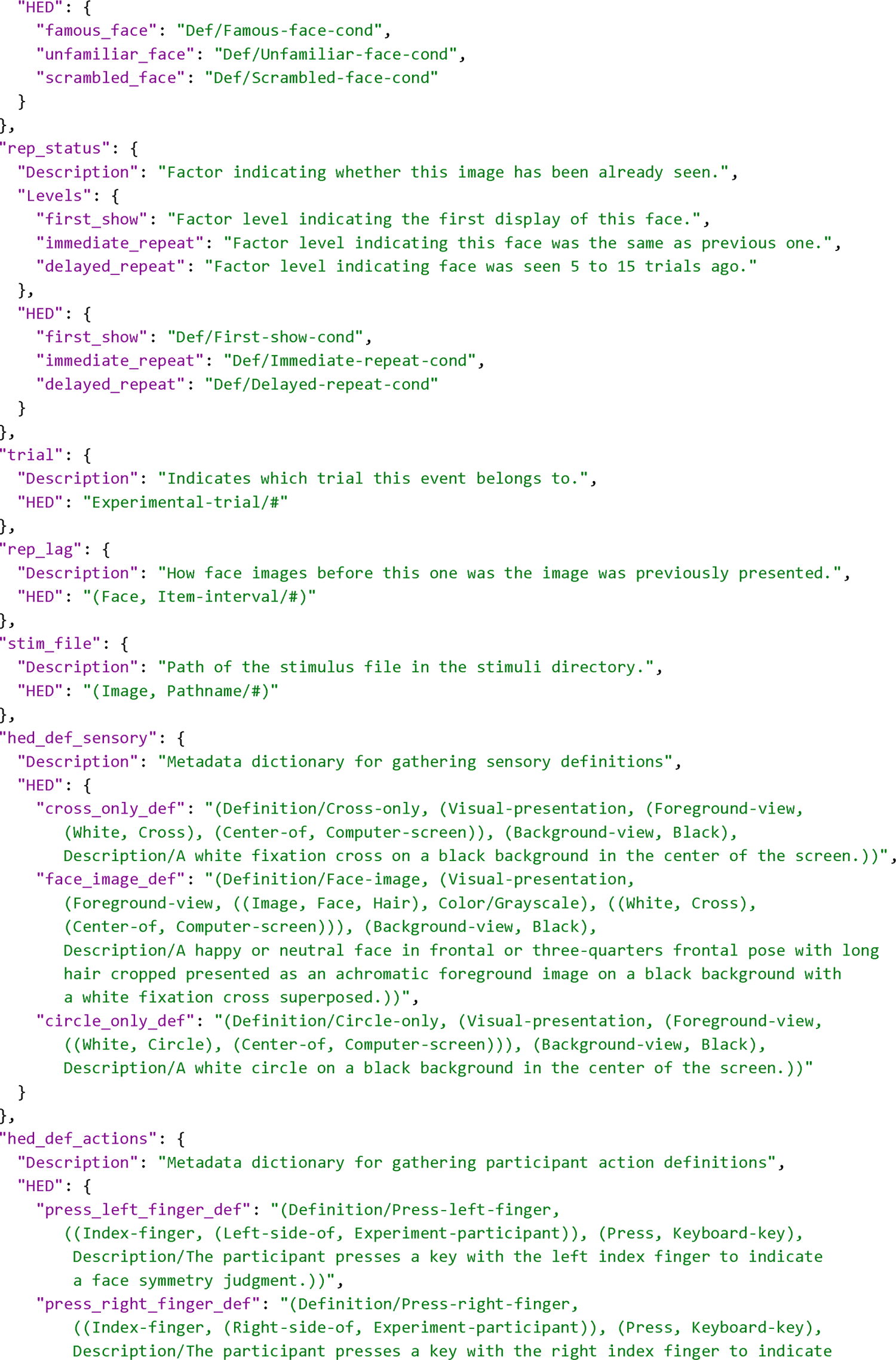
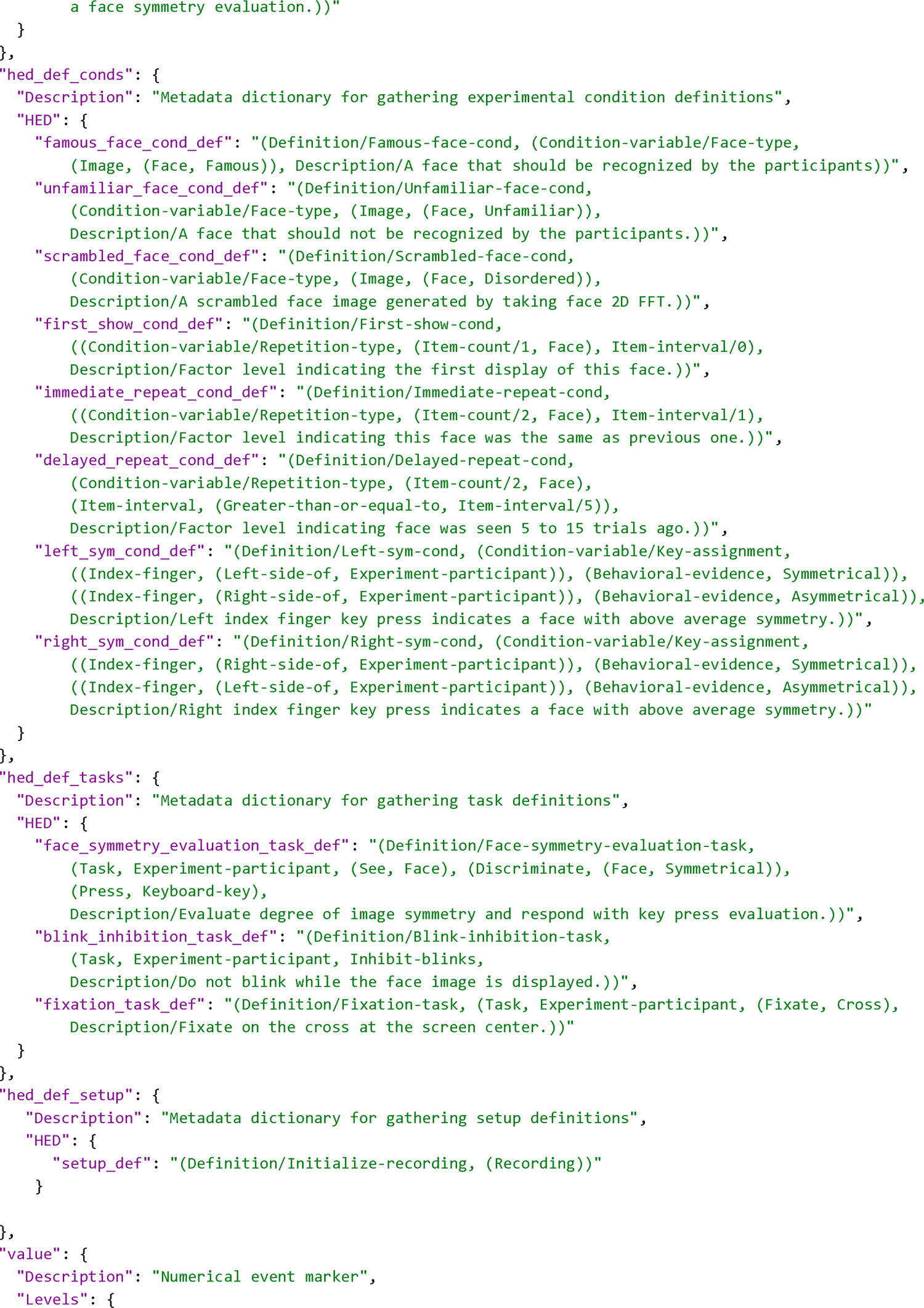
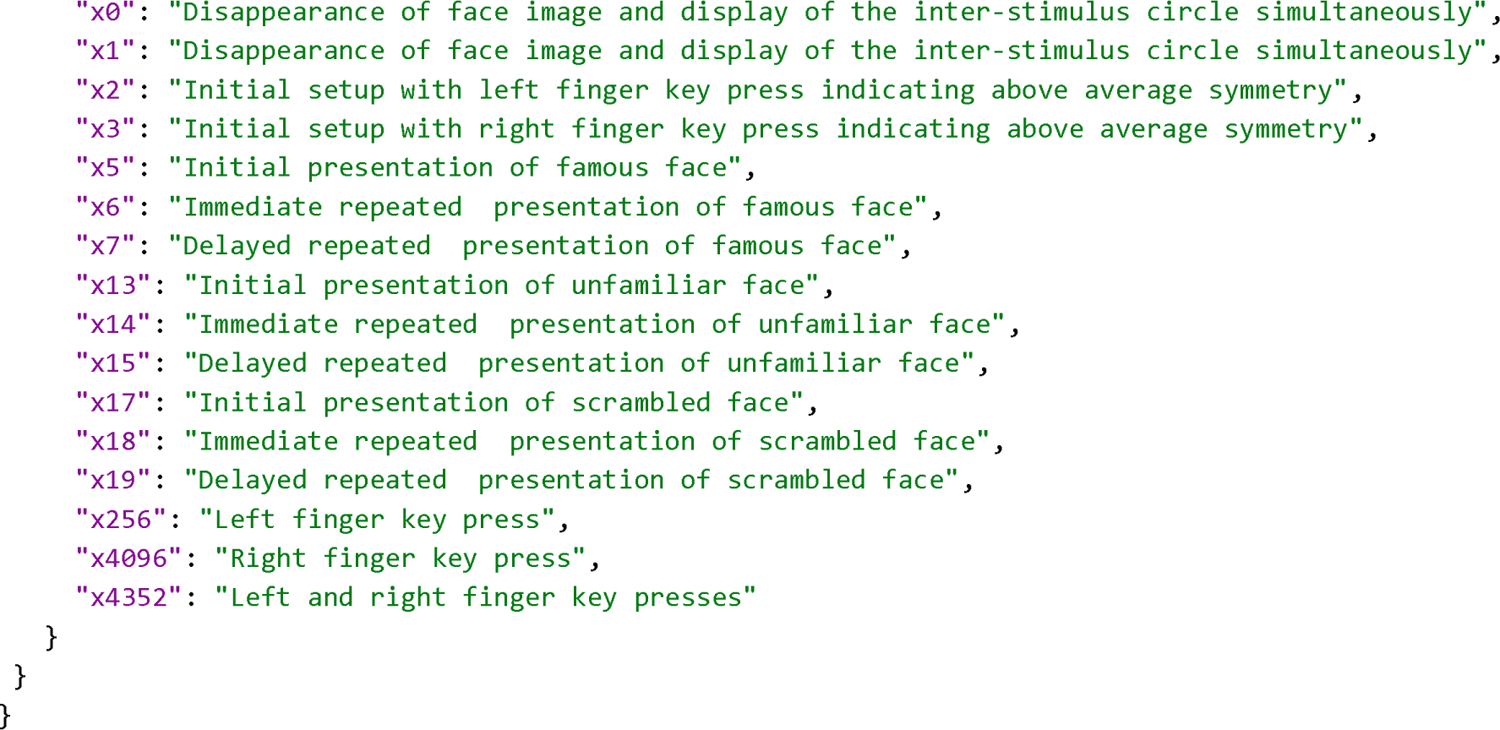
Complete JSON event file for Wakeman-Henson dataset (short-form). The file has been re-spaced for readability. All of the definitions have been gathered into additional metadata dictionaries at the end of the file. We have included the text descriptions as BIDS Levels for categorical columns, but these could also be included in the HED annotations using the *Description* tag.

**Supplementary Table 2:** The assembled form of the HED annotation for the second event in Table 3 (as shown in Table 5) and in three different forms expanded by tools. Form 1 is the form that would normally appear in the `…events.json` sidecar and be viewed. The tag strings have been re-spaced and partially bolded for readability.

**Form 1:**
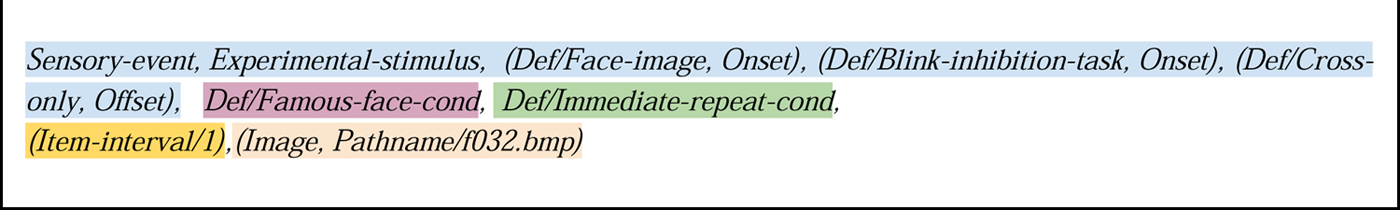
Short-form annotation of the sensory event corresponding to the first showing of famous face image f032.bmp. Definitions are unexpanded (as shown in Table 7).

**Form 2:**
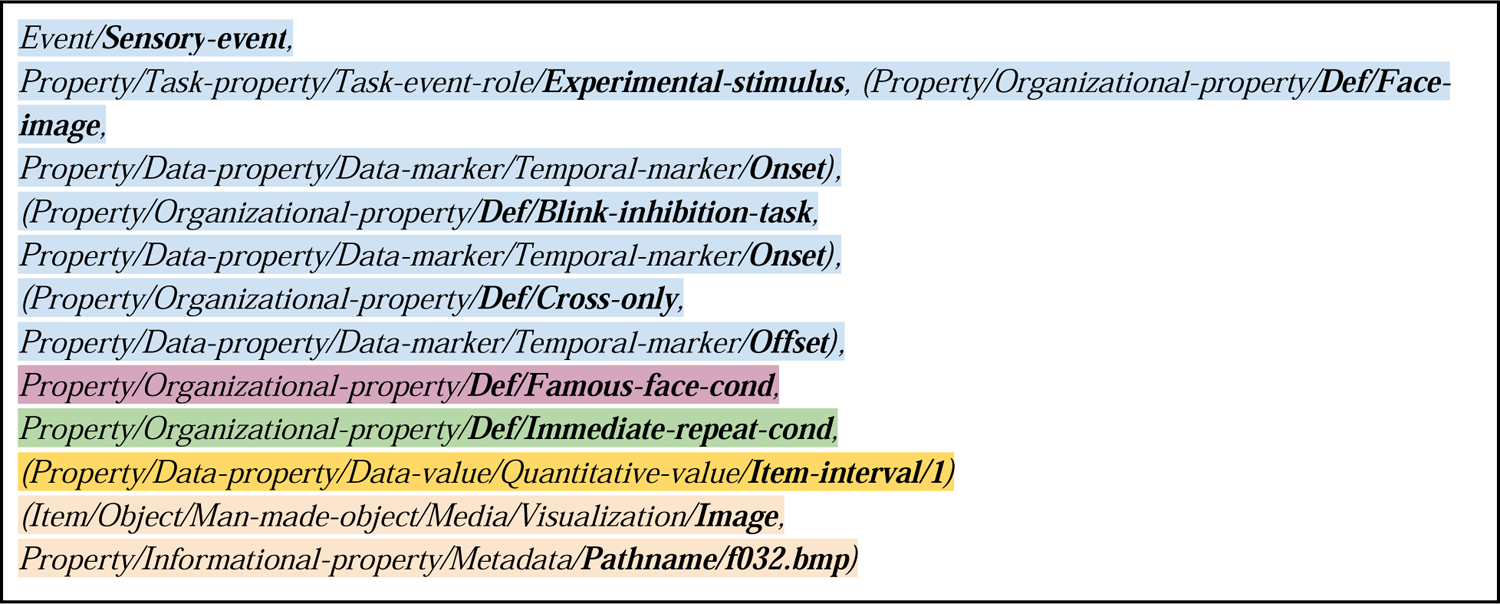
Long-form annotation of the sensory event corresponding to an immediate reshowing of famous face image f032.bmp. Definitions are unexpanded. Terms from Form 1 are shown in bold.

**Form 3:**
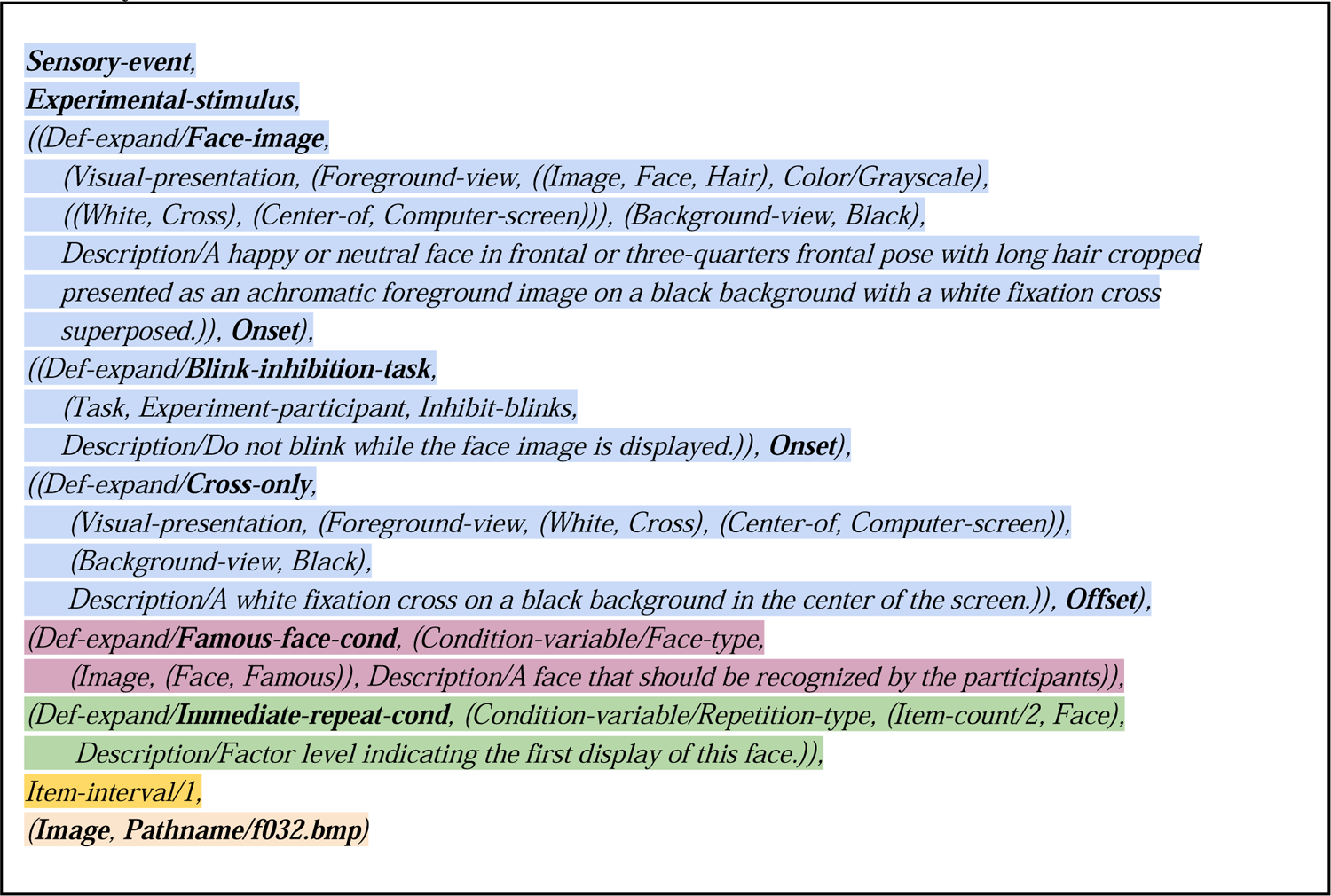
Short-form annotation of the sensory event corresponding to the immediate reshowing famous face image f032.bmp. Definitions are expanded. The annotation has been manually indented to improve readability.

**Form 4:**
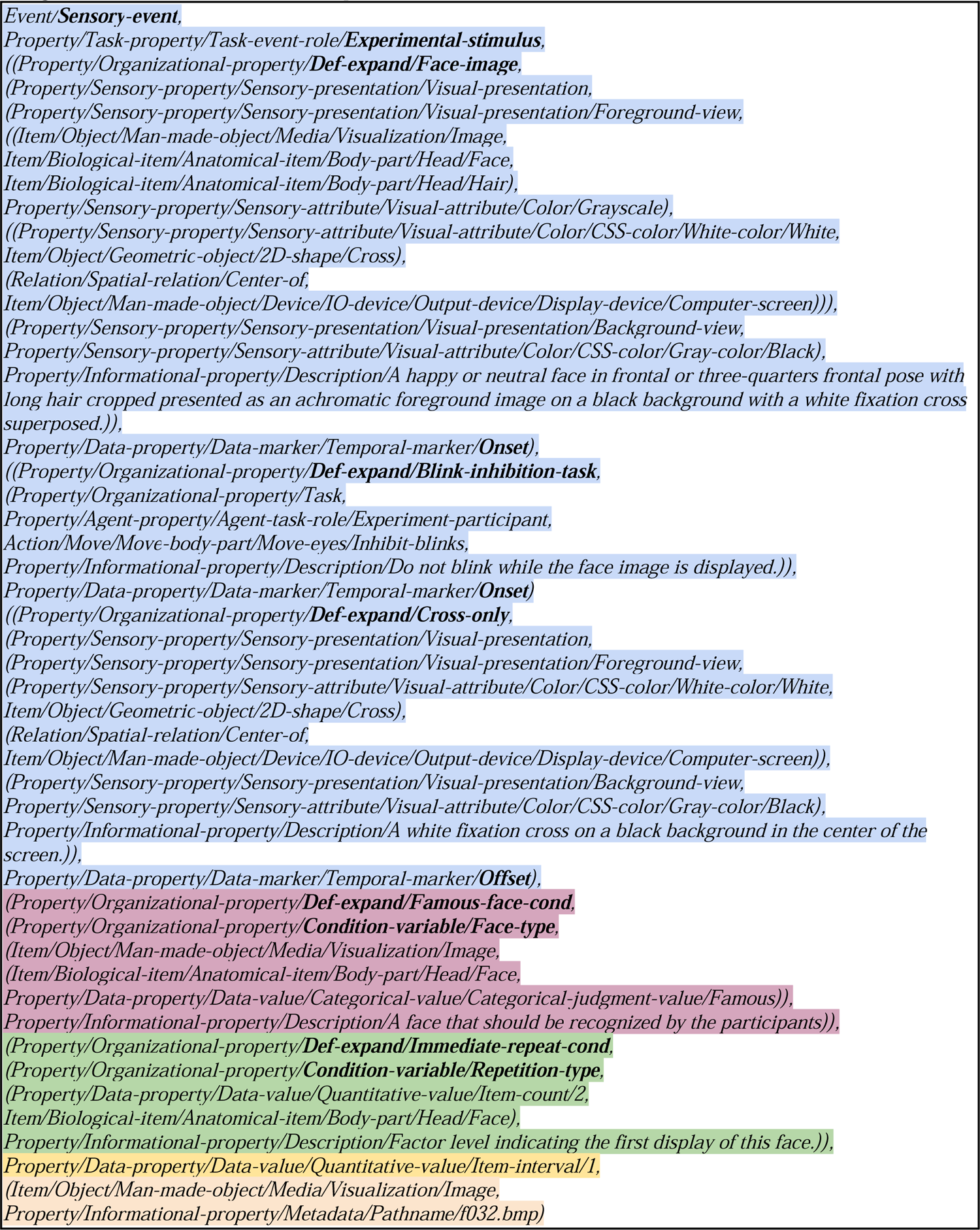
Long-form annotation of the sensory event corresponding to the first showing of famous face <u>image f032.bmp. Definitions are expanded.</u>

